# Direct Readout of Multivalent Chromatin Reader-Nucleosome Interactions by Nucleosome Mass Spectrometry

**DOI:** 10.1101/2025.05.01.651740

**Authors:** Alexander S. Lee, Nickolas P. Fisher, Matthew R. Marunde, Pei Su, Laiba F. Khan, Bria Graham, Hailey F. Taylor, Ugochi C. Onuoha, Taojunfeng Su, Kevin Jooß, Luis F. Schachner, Harrison A. Fuchs, Kelsey Noll, Marcus A. Cheek, Jonathan M. Burg, Zu-Wen Sun, Catherine A. Musselman, Michael-Christopher Keogh, Neil L. Kelleher

**Author notes:** **Correspondence:** Prof. Dr. Neil Kelleher [ ]. **Author Contributions**: A.S.L performed BPTF binding experiments with PTM-defined semi-synthetic nucleosomes using Nuc-MS. P.S., K.J., and L.F.S. assisted in MS analysis of BPTF binding experiments. A.S.L. performed affinity purification experiments. A.S.L, N.P.F., and T.S. acquired and analyzed Nuc-MS of CAP-endogenous nucleosome complexes. M.R.M., L.F.K., B.G., H.F.T. and U.C.O., M.A.C., K.N., J.M.B. and Z-.W.S. variously contributed to the synthesis of fully defined histones and nucleosomes; CAP construct design purification and characterization; and dCypher-Luminex experiments. H.A.F. assisted in purification of BPTF. M.-C.K. and C.A.M. supervised project performance and supported data analysis / interpretation. A.S.L. and N.L.K. conceived the study and drafted the initial manuscript. All authors contributed to and are responsible for subsequent versions. **Abbreviations:** Ammonium acetate (AmAc), Bromo adjacent homology (BAH), Bromodomain (BD), Bromodomain-containing protein 4 (BRD4), Bromodomain and PHD finger-containing transcription factor (BPTF), Chromatin-associated protein (CAP), native Mass Spectrometry (nMS), Native Top-Down Mass Spectrometry (nTDMS), Nucleosome Mass Spectrometry (Nuc-MS), Miccrococal nuclease (MNase), Glutathione S-transferase (GST), Chromodomain (CD), Plant homeodomain (PHD), *Populus trichocarpa* Short Half-life (PtSHL), post-translational modification (PTM).

## Abstract

Histone post-translational modifications (PTMs) often serve as distinct recognition sites for the recruitment of chromatin-associated proteins (CAPs) for epigenome regulation. While CAP-PTM interactions have been extensively studied using histone peptides, this cannot consider the regulatory potential of multi-site binding on intact nucleosomes. To overcome this limitation, we applied Nucleosome Mass Spectrometry (Nuc-MS), a native Top-Down MS approach that enables controlled disassembly of intact CAP:nucleosome (CAP:nuc) complexes to provide a direct readout of the contained histone proteoforms. As proof of principle, we show the BPTF PHD-BD native tandem reader requires coincident H3K4me3K9acK14acK18ac for effective nucleosome engagement. We extend our approach to explore how the BRD4 (native BD1-BD2), DNMT3A-MPP8 (chimeric PWWP-CD), and PtSHL (native BAH-BD) tandem readers interact with endogenous nucleosomes. Each reveals distinct enrichment profiles: BRD4 favoring di- and tri-acetylated histone H4 proteoforms, whereas DNMT3A-MPP8 and PtSHL preferentially interact with hypermethylated H3 proteoforms. Of note the latter enriches combinatorial {H3K4me3K27me3} on the same histone tail in HeLa chromatin, and thus expands the potential biology of this widely studied bivalent signature. By directly characterizing CAP:nuc complex composition with combinatorial PTM information in a single readout, Nuc-MS serves as a new approach to discover the modifications driving binding, and therefore primary candidates to explore for structural biology and genomic studies.

For TOC only.
Synopsis.
Nuc-MS provides the ability to control the disassembly of chromatin-associated protein-nucleosome complexes (CAP-nuc) and delineate the histone proteoforms driving tandem reader domain binding.

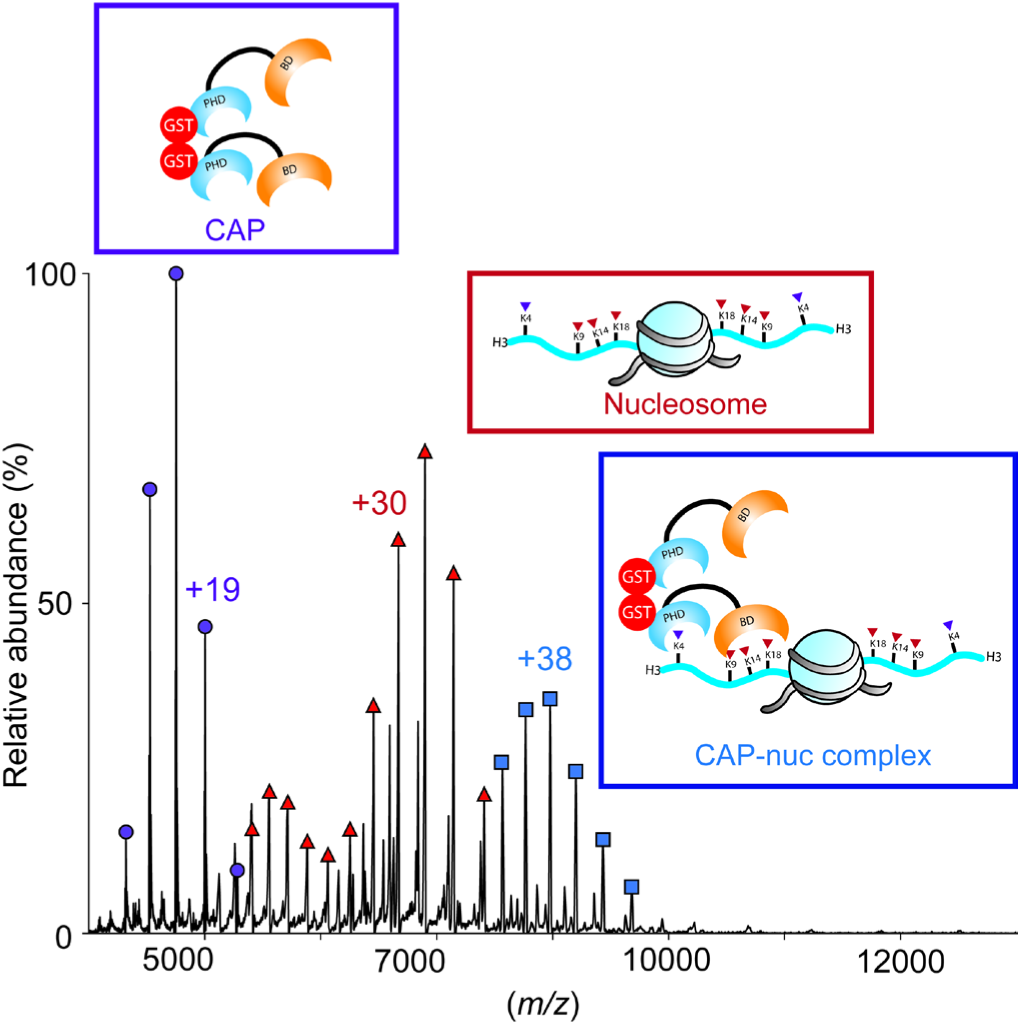

## 1. Introduction

Nucleosomes serve as the fundamental repeating units of eukaryotic genome organization, with arrays of them forming higher-order chromatin.^1^ Canonical nucleosomes comprise two copies of each core histones (H2A, H2B, H3 and H4), forming protein octamers wrapped by 147 bp of DNA.^1^ Within this unit, the globular histone-fold domains mediate extensive structural associations with each other and DNA, while highly charged N-terminal tails dynamically interact with various intra- and inter-nucleosomal surfaces.^1–3^ Each histone can be decorated with myriad post-translational modifications (PTMs), added and removed by a diversity of histone-modifying enzymes.^4^ Within chromatin, local nucleosome context (the nucleoform^5–7)^ is comprised of the encompassed histones, their variants and PTMs (*aka* the included proteoforms).^8^ To mediate function, PTMs (or more accurately PTM combinations) serve as recognition sites for chromatin-associated proteins (CAPs) through specialized reader domains.^9^ The regulated recruitment of CAPs to nucleosomes at a given genomic location supports the assembly of larger nucleoprotein complexes that govern DNA-centric processes including DNA replication, transcription, and repair.^10,11^

For decades, most studies that interrogate the binding potential between CAPs and PTMs have used histone peptide pulldowns or arrays, predictions from structures of related reader domains with histone peptides, or domain incubations with cell extracts followed by immunoblotting with PTM-specific antibodies.^12–19^ Such approaches are often undermined by the reductive formats (minimal reader domains and histone peptides) that poorly represent the regulatory potential of a CAP:nucleosome (CAP:nuc) complex, and utilize anti-PTM antibodies of varying quality.^14,17,20^ Using such approaches, histone peptides can inform on CAP engagement to combinatorial PTMs on the same histone tail (in *cis*), but not on interactions across nucleosomal histones (in *trans*).^21^ While histone N-terminal tails are traditionally depicted as extending from the nucleosome core, current studies show they instead dynamically associate with nucleosomal DNA with profound regulatory potential.^22–25^ Histone peptides do not supply the nucleosomal DNA engaged by readers that recognize a PTM in this context, such as PWWP domains or certain bromodomains.^23,26–30^ Lastly, histone peptides cannot provide the diversity of ‘non-specific’ surfaces that support multivalent CAP:nuc engagement, such as the wrapping of DNA and the acidic patch.^10,19,31–35^ Semi-synthetic PTM-defined nucleosomes provide the ability to interrogate CAP interactions with targets more representative of their natural state, but require direct chemical synthesis of each site-specific PTM and their assembly into combinatorial nucleosomes. As such, it is cost-prohibitive to address the immense potential diversity of histone proteoforms in chromatin.^36–39^ Rather, an approach is needed to screen for the binding of CAPs (alone or within megadalton complexes) to nucleosomes absent *a priori* knowledge of preference, and then detect the various enriched histone proteoforms and nucleoforms for further study.

To this end, MS-based proteomics has contributed a lot to our understanding CAP:nuc interactions.^40–47^ However, current MS approaches often require the nucleosomes and any associated CAPs be denatured and digested to short peptides prior to analysis (*aka*. bottom-up proteomics).^40–42^ This loses information on CAP complex composition and histone PTM co-occurrence (unless immediately adjacent). Due to this, we still have a limited understanding of how most histone proteoforms contribute to epigenetic regulation, cellular identity, and disease development. Advances in MS can inform on complete histone proteoforms, including variants and distal PTM combinations (*e.g.*, {H3.2K36me2K79me2}).^5,6,48–52^ Native Top-Down Mass Spectrometry (nTDMS) refers to the controlled disassembly of protein complexes and subsequent characterization of encompassed proteoforms.^53,54^ To provide a new approach, we utilized Nucleosome Mass Spectrometry (Nuc-MS) to inform on the diverse histone proteoforms that compose either semi-synthetic [fully defined] or endogenous {native} nucleosomes (*aka* the nucleoform) in CAP:nuc complexes.^5–7^

Here, we broaden the application of Nuc-MS to: **i)** confirm the PTMs required for CAP interaction with fully defined nucleoforms; and **ii)** identify the histone proteoform landscape after CAP-mediated selection from a pool of endogenous nucleoforms. As proof of principle, we confirm the preference of the BPTF PHD-BD tandem reader for PTM-defined ([H3K4me3K9acK14acK18ac]) nucleosomes^25^, and that individual loss-of-function mutations in either reader domain prevents stable complex formation. We then explore the binding of three reader-domain tandems with endogenous nucleosomes. Here a tandem bromodomain (BD1-BD2) from BRD4 enriches nucleosomes containing acetylated H2A.Z and histone H4 proteoforms, including one with a rare acetylation at H4K44. While the DNMT3A-MPP8 chimeric PWWP-CD and PtSHL native BAH-BD recover ({H3K9me3K36me3}) and ({H3K4me3K27me3K79me2}), respectively. These results highlight the potential of Nuc-MS as a tool to discover the combined elements driving CAP:nuc interactions, and allow for focused subsequent studies and improve mechanistic insight into epigenetic regulation.

## 2. Results and Discussion

The Nuc-MS workflow provides three MS-based elements to interrogate CAP:nuc interactions: **i)** intact mass analysis of native CAP:nuc complexes (MS1); **ii)** intact mass analysis of histone subunits ejected from CAP:nuc complexes (MS2); and, **iii)** tandem MS fragmentation data to assert the enriched histone proteoforms (MS3) **(Figure 1)**.

**Figure 1:**
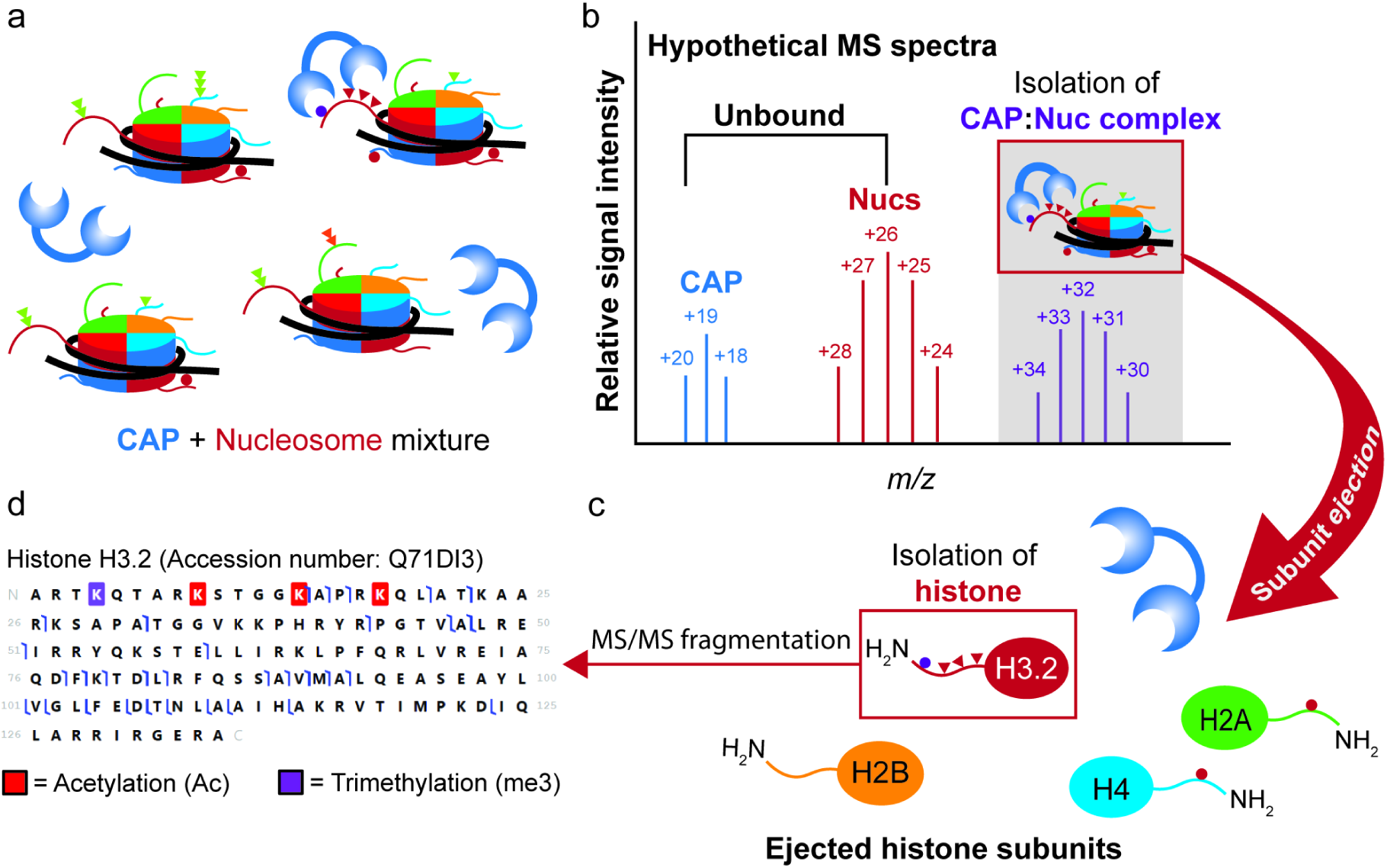
Nucleosome Mass Spectrometry (Nuc-MS) workflow for the direct analysis of CAP-associated nucleoforms. **a)** Chromatin-associated proteins (CAPs) are mixed with Nucleosomes (semi-synthetic or endogenous). **b)** Native and intact CAP-nucleosome complexes are isolated (MS1). **c)** These complexes are activated by collision with nitrogen gas to eject histone subunits (MS2). **d)** The desired histone subunit is further isolated and fragmented (pseudo MS3) to determine protein sequence and PTM identity and position.

To begin, we conducted a Luminex-based assay to confirm the binding preference of GST-BPTF PHD-BD (hereafter BPTF) binding preference for PTM-defined histone H3 peptides and semi-synthetic nucleosomes **(SI Figure 1)**.^55^ As expected, BPTF effectively bound H3_[1-20]_K4me3 and H3_[1-20]_K4me3tri^Ac^ (K4me3K9acK14acK18ac) peptides (with respective EC_50_^Rel^ values of 4.30 and 18.9 nM: **SI Figure 1B** and **SI Table 1A-B)**. However, in the nucleosome context, BPTF showed a strong preference for the ([H3K4me3tri^Ac^]) combinatorial (EC_50_^Rel^ value of 23.4 nM: **SI Figure 1C** and **SI Table 1C-D)**.^25,56^

We next used Nuc-MS (MS1) to examine this CAP:nuc interaction, and determined that BPTF failed to stably engage ([unmodified]), ([H3K4me3]) or ([K4acK9acK14acK18ac]) (H3 tetra^Ac^) nucleosomes. This was in strong contrast with the binding of BPTF to the nucleoform containing three acetylations with H3K4me3 ([H3K4me3tri^Ac^]) **(Figure 2A-D)**.^25^ Further, loss-of-function alanine mutations^16,57^ within the BPTF PHD finger (W2891A, PHD*), bromodomain (N3007A, BD*), or both domains (PHD*BD*) abrogated complexation with [H3K4me3tri^Ac^] (**SI Figure 3** and **SI Table S1**). Through disruption of the stable BPTF:nucleosome complex inside the mass spectrometer (MS2), we validated the identity of each histone proteoform at the intact level, and their PTMs by another state of fragmentation in the gas phase (MS3) **(SI Figure 2)**. Of note, this complex contained two copies of BPTF, though we were unable to discern if this was two independent bindings, or one binding dimerized via the N-terminal GST-tag.^58^

**Figure 2:**
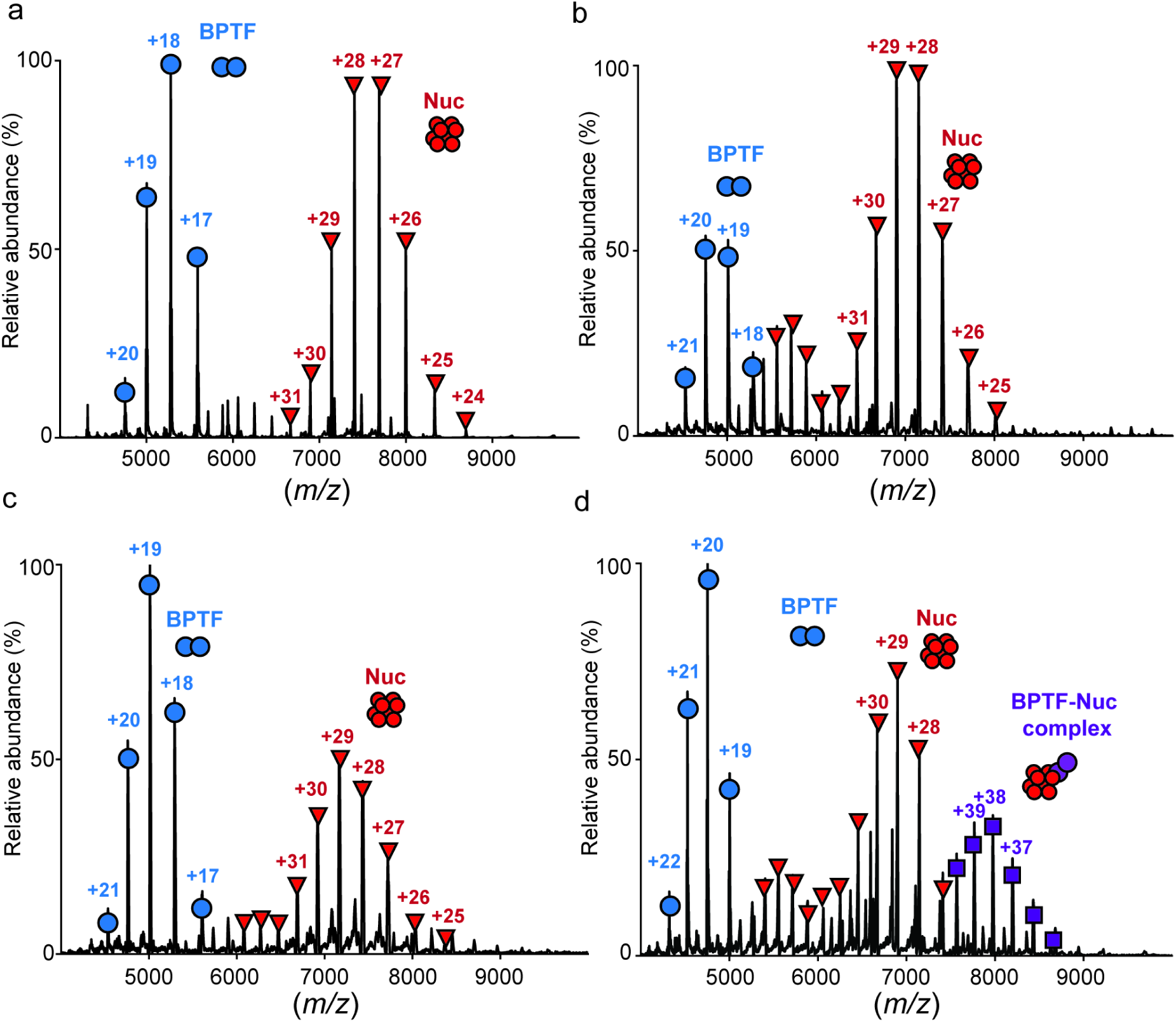
Nuc-MS analysis of BPTF PHD-BD binding to PTM-defined semi-synthetic nucleoforms. **a-d)** Native MS1 spectrum of intact GST-BPTF-PHD-BD with ([unmodified]) **(a)**, ([H3K4me3]) **(b)**, ([H3tetra^Ac^]) **(c)**, or ([H3K4me3tri^Ac^]) **(d)** mononucleosomes. The absence of complexation in **(a)** is reflected by charge state distributions at (+17 to +20) and (+24 to +38), respectively corresponding to a BPTF dimer (possibly via the N-terminal GST-tag^58^) and unmodified nucleosome. Complexation with ([H3K4me3tri^Ac^]) in **(d)** is reflected by the appearance of a charge state at (+35 to +41). Representative of three independent experiments shown.

In the above, our Nuc-MS approach can successfully detect multivalent CAP binding to fully defined semi-synthetic nucleosomes and delineate the histone proteoforms necessary for protein complexation. We next challenged the approach to characterize CAP-enriched endogenous nucleosomes derived from HeLa cells, hypothesizing that different tandem reader domains should recover distinct histone proteoform landscapes. To achieve this, we mixed each CAP with a pool of endogenous nucleosomes liberated from HeLa chromatin by Micrococcal nuclease (MNase) digestion (see **Experimental Methods and Materials** and **SI Figures 4** and **5**).

Bromodomain-containing protein 4 (BRD4) is a member of the Bromodomain and Extra-Terminal (BET) family, and contains tandem bromodomains (BD1 and BD2) to mediate interaction with nucleosomes hyperacetylated at histones H3 and H4.^26,59–65^ BRD4 mediates roles in DNA repair, higher order chromatin structure, and transcription^61,64,66–72^, with dysfunction associated with a range of cancer subtypes^73,74^, and BRD4-targeting therapeutics in active development.^75–82^ The 6xHis-BRD4 BD1-BD2 tandem reader (hereafter BRD4) **(SI Figure 6)** was found to enrich for hyperacetylated H4 proteoforms, with 0.5, 2.6 and 4.5-fold increases in the amount of mono-, di-, and tri-acetylated forms relative to bulk HeLa nucleosomes **(Figure 3A)**. Tandem MS characterization of these H4 proteoforms revealed multiple acetylations at K5, K8, K12, and K16 (reflective of active transcription and euchromatin^26,59–65,83^) in combination with K20me2 **(Figure 3A** and **SI Figure 7)**. We additionally identified the presence of a new {H4K20me2K44ac} proteoform **(Figure 3C)** not previously linked to canonical BRD4 function but may reflect a role in DNA damage and repair.^69,70,84,85^

**Figure 3:**
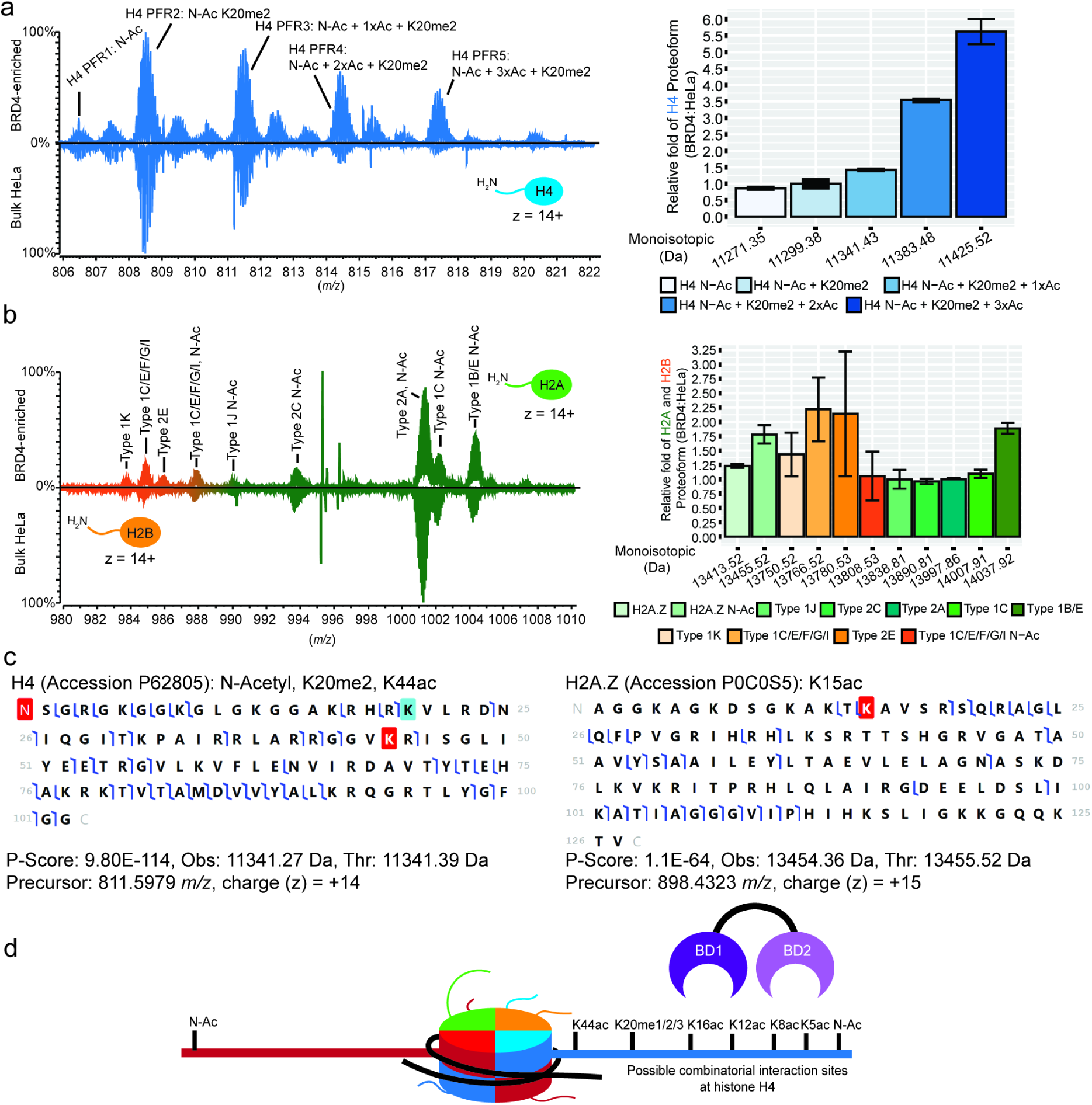
Nuc-MS analysis of BRD4-enriched endogenous nucleoforms. **a-b)** Representative MS1 spectra of the histone H4 **(a)** and H2A **(b)** proteoform landscapes (left) and their quantification (right)) after nucleosome enrichment by the native tandem reader BRD4 BD1-BD2. In each landscape, acetyl equivalents are represented as (nxAc), with BRD4-enriched nucleosomes above and bulk HeLa nucleosomes below. **c)** Graphical fragment maps of H4: N-Acetyl, K20me2, K44ac (left) and H2A.Z: K15ac (right). **d)** Model of possible BRD4 PTM-binding sites in nucleosome context. Quantification of enrichment was calculated by determining the relative abundances of each histone proteoform and normalizing to either H4: N-Acetyl + K20me2 or Histone H2A Type 2A. The relative ratio of BRD4:HeLa bulk is the normalized values of each histone proteoform enriched by BRD4 divided by that in bulk HeLa. Error bars represent standard deviation from the mean.

On further analysis of the BRD4:nuc complex, proteoforms of histone H3.2 were consistently below the level required for confident tandem MS characterization (not shown), though all that crossed the detection threshold were unchanged relative to bulk HeLa nucleosomes **(Figure 3A** and **SI Figure 8)**. Examination of the enriched histone H2A and H2B proteoforms identified no major differences relative to bulk **(Figure 3B** and **SI Figure 9)**, with the exception of 75% enrichment for an acetylated form of H2A.Z {H2A.ZK15ac}, a variant often associated with gene transcription in development and differentiation **(Figure 3C** and **SI Figure 10)**.^68,86–88^ Of note, this insight regarding to BRD4 preferred nucleoforms, for example ({H2A.ZK15ac}•{H4K5acK12acK16acK20me2K44ac}) was not previously identified by peptide-centric MS and other biochemical approaches.

We next expanded our study to the histone proteoform landscapes enriched by tandem reader domains that engage lysine methyl states. The DNMT3A-MPP8 reader fuses the H3K36me2/3 binding PWWP domain from *de novo* DNA methyltransferase 3A to the H3K9me3 binding chromodomain of M-phase Phosphoprotein 8 (MPP8) **(SI Figure 11)**. This chimera has been shown to be an effective tool for genomic and biochemical studies^89,90^, and its combinatorial PTM targets a bivalent state (‘repressive’ H3K9me3 co-incident with ‘transcriptionally active’ H3K36me3) marking poised transcriptional enhancers.^91^ Nuc-MS of CAP:nuc complexes assembled from DNMT3A-MPP8 and endogenous HeLa nucleosomes (**SI Figure 4**) identified a proteoform landscape **>**50% enriched for histone H3.2 containing three or more methyl equivalents **(Figure 4A)**. Isolation and subsequent tandem MS fragmentation (MS2) of these histone H3 proteoforms identified the expected {H3.2K9me3H3K36me3} **(Figure 4A** and **SI Figure 12)**.^89,90^ Characterization of the other histone proteoforms revealed enrichment of {H4K16acK20me2}, both PTMs associated with active transcription **(Figure 4B** and **SI Figure 13)**^50,51,92^, but no obvious change in H2A or H2B proteoforms relative to bulk HeLa nucleosomes **(Figure 4C** and **SI Figure 14**). These findings **(Figure 4D)** are in line with prior orthogonal studies ^89,90^, and validate the sensitivity of Nuc-MS as an approach to usefully interrogate CAP:endogenous nuc complexes.

**Figure 4:**
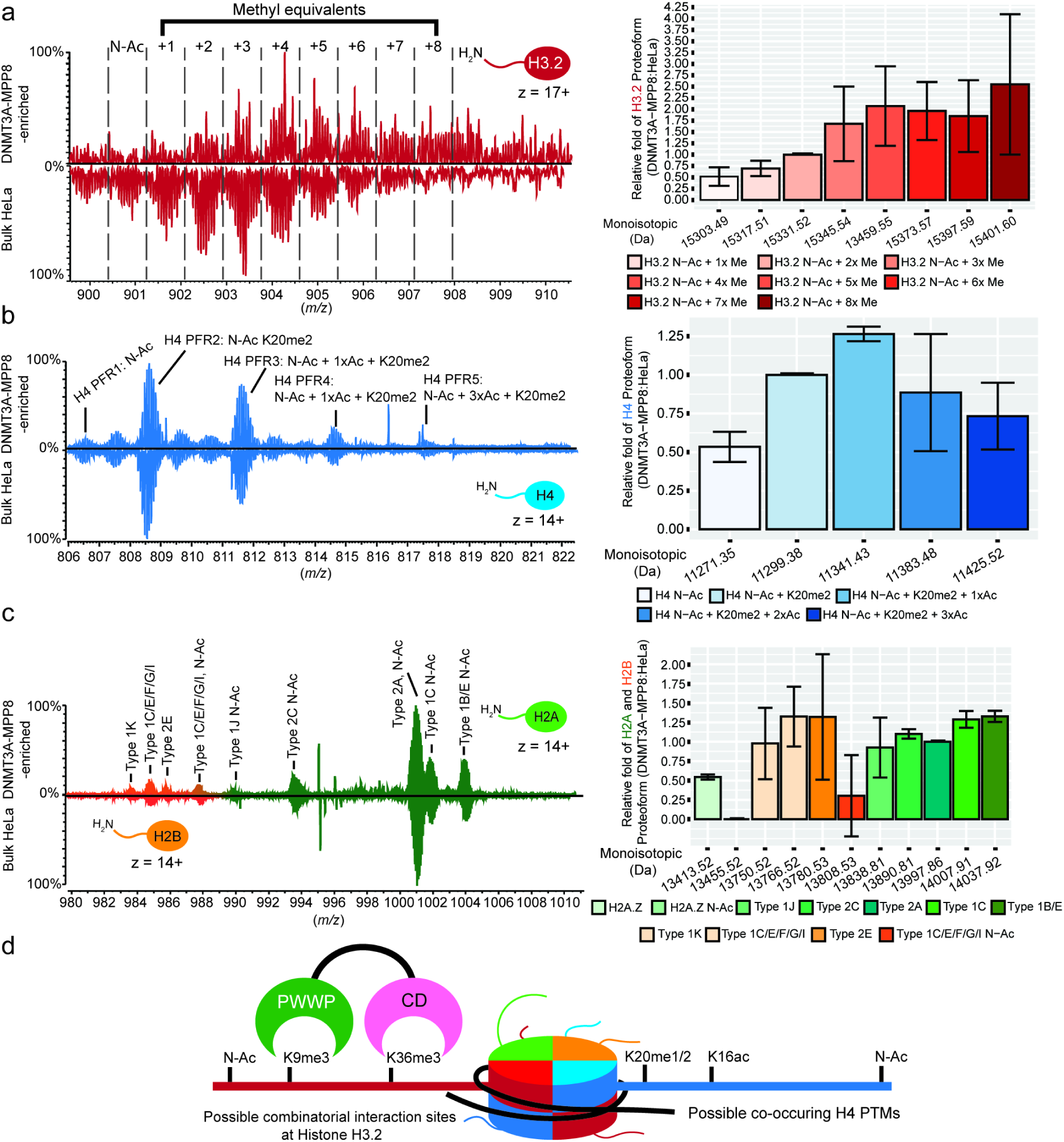
Nuc-MS analysis of DNMT3A-MPP8-enriched endogenous nucleoforms. **a-c)** Representative MS1 spectra of the histone H3 **(a)**, H4 **(b)** and H2A/H2B **(c)** proteoform landscapes (left) and their quantification (right) after nucleosome enrichment by the chimeric tandem reader DNMT3A-MPP8 PWWP-CD. In each landscape acetyl and methyl equivalents are respectively represented as (nxAc) or (nxMe), with DNMT3A-MPP8 nucleosomes above and bulk HeLa mononucleosomes below. **d)** Model of possible DNMT3A-MPP8 PTM-binding sites in nucleosome context. Quantification of enrichment was calculated by determining the relative abundances of each histone proteoform and normalizing to either H3.2: N-Ac + 3xMe, H4: N-Acetyl + K20me2, or Histone H2A Type 2A. The relative ratio of DNMT3A-MPP8:HeLa bulk is the normalized values of each histone proteoform enriched by DNMT3A-MPP8 divided by that in bulk HeLa. Error bars represent standard deviation from the mean.

In the final example, Short Half Life (PtSHL) from the plant *Populus trichocarpa* contains a bromodomain-adjacent homology (BAH) domain and PHD finger reader in tandem that **(SI Figure 15)** recognizes H3K4me3 and H3K27me3, with this binding important for SHL-mediated floral repression.^93^ In mammalian cells H3K4me3 and H3K27me3 are usually mutually exclusive, respectively marking transcriptionally active promoters and polycomb repressed regions. An interesting exception is ‘bivalency’, where the functionally opposing PTMs co-exist within the same nucleosome but on distinct histone H3 tails (so a heterotypic state), in embryonic stem cells, potentially to poise genes for rapid active or repression during development.^46,94–96^

Nuc-MS of CAP:nuc complexes assembled from PtSHL and endogenous HeLa nucleosomes revealed a histone H3.2 proteoform profile with increased methyl equivalents, while the H2A, H2B and H4 (largely {H4K20me1/2} or {H4K16acK20me1/2}) proteoform profiles were similar to bulk HeLa **(Figure 5**, and **SI Figures 4**, **16-18)**. We conducted tandem MS fragmentation of the H3 proteoforms with three or more methylation equivalents and identified {H3.2K4me3K27me3K79me2} **(Figure 5A** and **SI Figure 16)**. The specific enrichment of this *cis* proteoform supports the functionality of each domain in the PtSHL BAH-PHD tandem reader and suggests a major dysregulation of the canonical bivalent mark in HeLa cells.^97,98^

**Figure 5:**
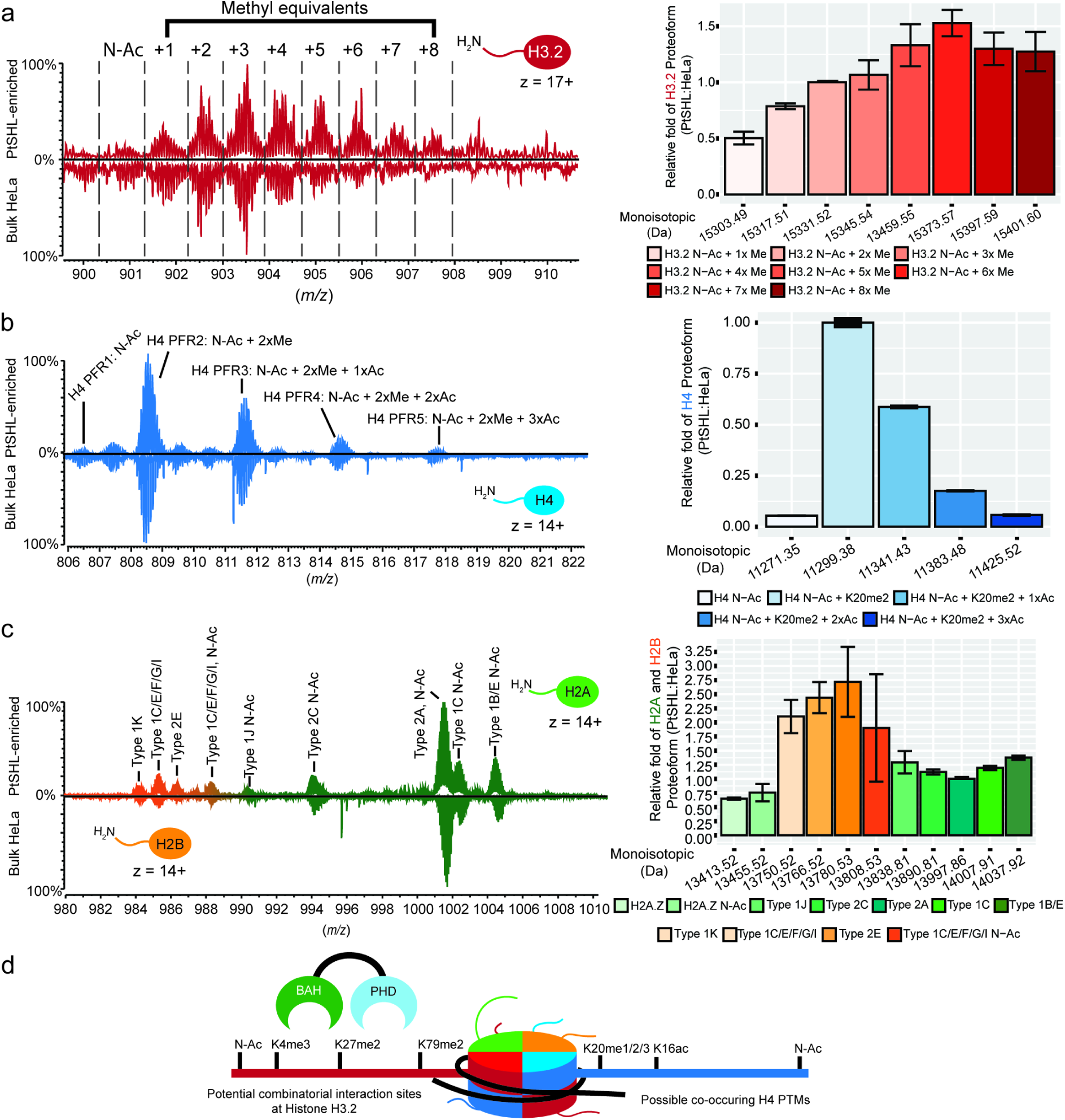
Nuc-MS analysis of PtSHL-enriched endogenous nucleoforms. **a-c)** Representative MS1 spectra of the histone H3.2 **(a)**, H4 **(b)** and H2A/H2B **(c)** proteoform landscapes (left) and their quantification (right) after nucleosome enrichment by the native tandem reader PtSHL BAH-PHD. In each landscape acetyl and methyl equivalents are respectively represented as (nxAc) or (nxMe), with PtSHL-enriched above and bulk HeLa nucleosomes below. **d)** Model of possible PtSHL PTM-binding sites in nucleosome context. Quantification of enrichment was calculated by determining the relative abundance of each histone proteoform and normalizing to either H3.2: N-Ac + 3xMe, H4: N-Acetyl + K20me2, or Histone H2A Type 2A. The relative ratio of PtSHL:HeLa bulk is the normalized values of each histone proteoform enriched by PtSHL divided by that in bulk HeLa. Error bars represent standard deviation from the mean.

## 3. Conclusion

Directly informing on the functional relationship between CAPs and the chromatin landscape is a profound challenge for existing methods^7^, which are often unable to dissect the relative contribution of diverse histone proteoforms to CAP engagement. To address this, we applied Nuc-MS to investigate the proteoform landscape within four tandem-reader CAP:nuc complexes. This includes proof of principle with fully defined nucleosomes (BPTF PHD-BD: ([H3K4me4tri^Ac^])); an exploration of capability relative to prior biochemical and genomic approaches with endogenous nucleosomes (BRD4 BD1-BD2: ({H2A.ZK15ac}•{H4K5acK12acK16ac K20me2K44ac}) and DNMT3A-MPP8 PWWP-CD: ({H3.2K9me3 H3K36me3}); and a new interrogation of PtSHL BAH-PHD: ({H3.2K4me3K27me3K79me2}) with endogenous nucleosomes (**Figures 2-5** and **SI Figures 1-18**). In each case we validated known interactions, but also revealed novel histone proteoform compositions showing the potential of our new approach to focus downstream studies such as ChIP-seq to specific histone modifications. Its further development and application to multi-subunit CAP complexes and chromatin from healthy and disease states should offer insights to the importance of epigenetic (dys)regulation.

## Supporting Information

Supporting information includes experimental methods, materials, and additional references pertaining to this study. **SI Figure 1**: Differential BPTF binding preference for histone peptides and nucleosomes; **SI Figure 2**: Nuc-MS characterization of BPTF-bound ([H3K4me3tri^Ac^]) nucleosomes; **SI Figure 3**: Both domains in the BPTF-PHD-BD tandem are required for effective binding to ([H3K4me3tri^Ac^]) nucleosomes; **SI Figure 4-5**: Purification and validation of CAP-endogenous nucleosomes; **SI Figure 6**: Characterization of 6xHis-BRD4-BD1-BD2 native tandem reader; **SI Figure 7:** Characterization of Histone H4 proteoforms from BRD4-enriched endogenous nucleosomes; **SI Figure 8**: Relative quantification of H3.2 proteoforms from BRD4-enriched endogenous nucleosomes; **SI Figures 9**: Characterization of Histone H2A and H2B proteoforms from BRD4-enriched endogenous nucleosomes; **SI Figure 10**: BRD4 enriches for acetylated histone H2A.Z variant; **SI Figure 11**: Characterization of GST-DNMT3A-MPP8 PWWP-CD chimeric tandem reader; **SI Figures 12-14**: Tandem MS characterization of histone proteoforms from DNMT3A-MPP8-enriched endogenous nucleosomes; **SI Figure 15**: Characterization of GST-PtSHL-BAH-PHD native tandem reader; **SI Figure 16-18**: Tandem MS characterization of histone proteoforms from PtSHL-enriched endogenous nucleosomes; **SI Tables 1A-D**: Relative effective concentrations of binding affinities of BPTF and mutants with histone peptides and nucleosomes; **SI Tables 2-6**: MS instrumentation parameters used for characterization of histone proteoforms and CAPs; **SI Table 7**: MS instrumentation parameters used for tandem MS characterization of endogenous histone H3.2 proteoforms by Individual Ion Mass Spectrometry.

## Acknowledgements

This work was supported by the National Institutes of Health (NIH) National Institute of General Medical Sciences P41GM108569 for the National Resource for Translational and Developmental Proteomics at Northwestern University and NIA grant (RF1 AG063903). A.S.L is a trainee fellow under the Chemistry of Life Processes Predoctoral Training Grant (5T32GM105538-10) at Northwestern University. N.P.F is a trainee fellow under the Molecular Biophysics Predoctoral Training Grant (T32GM140995) at Northwestern University. *EpiCypher* was supported by NIH grants R44GM116584, R44GM136172, R44HG010595 and R43CA236474. Work in the Musselman laboratory is funded by NIH grant R35GM128705.

## Conflicts of interest

NLK serves as a consultant to Thermo Fisher Scientific and engages in entrepreneurship in the area of Top-Down Proteomics. *EpiCypher* is a commercial developer and supplier of reagents (*e.g.*, fully defined semi-synthetic nucleosomes) used in this study. M.R.M., L.K., B.G., H.F.T., U.C.O., K.N., M.A.C., J.M.B., Z.-W.S. and M.-C.K. own shares in *EpiCypher* with M.-C.K. also a board member of same.

