## Supplemental Information for "Direct Readout of Multivalent Chromatin Reader-Nucleosome Interactions by Nucleosome Mass Spectrometry"

### Table of Contents: Total 38 pages


- Experimental Methods and Materials
- Additional References

- **SI Figure 1:** Differential BPTF binding preference for histone peptides and nucleosomes.
- **SI Figure 2:** Nuc-MS characterization of BPTF-bound ([H3K4me3tri$^{Ac}$]) nucleosomes.
- **SI Figure 3:** Both domains in the BPTF-PHD-BD tandem are required for effective binding to ([H3K4me3tri$^{Ac}$]) nucleosomes.
- **SI Figure 4:** Extraction and affinity purification of CAPs complexed with endogenous nucleosomes.
- **SI Figure 5:** Purification of nucleosomal DNA from endogenous HeLa mononucleosomes.
- **SI Figure 6:** Characterization of 6xHis-BRD4-BD1-BD2 native tandem reader.
- **SI Figure 7:** Characterization of Histone H4 proteoforms from BRD4-enriched endogenous nucleosomes.
- **SI Figure 8:** Relative quantification of H3.2 proteoforms from BRD4-enriched endogenous nucleosomes.
- **SI Figure 9:** Characterization of Histone H2A and H2B proteoforms from BRD4-enriched endogenous nucleosomes.
- **SI Figure 10:** BRD4 enriches for acetylated histone H2A.Z variant.
- **SI Figure 11:** Characterization of GST-DNMT3A-MPP8 PWWP-CD chimeric tandem reader.
- **SI Figure 12:** Characterization of {H3.2 N-Acetyl, K9me3, K36me3} proteoform from DNMT3A-MPP8-enriched endogenous nucleosomes.
- **SI Figure 13:** Characterization of Histone H4 proteoforms from DNMT3A-MPP8-enriched endogenous nucleosomes.
- **SI Figure 14:** Characterization of Histone H2A and H2B proteoforms from DNMT3A-MPP8-enriched endogenous nucleosomes.
- **SI Figure 15:** Characterization of GST-PtSHL-BAH-PHD native tandem reader.
- **SI Figure 16:** Characterization of {H3.2 N-Acetyl, K4me3, K27me3, K79me2} proteoform from PtSHL-enriched endogenous nucleosomes.
- **SI Figure 17:** Characterization of Histone H4 proteoforms from PtSHL-enriched endogenous nucleosomes.
- **SI Figure 18:** Characterization of Histone H2A and H2B proteoforms from PtSHL-enriched endogenous nucleosomes.
- **SI Table 1A:** Relative Effective Concentration ($EC_{50}^{Rel}$) in Luminex assays to measure the interaction between GST-BPTF-PHD-BD constructs and PTM-defined histone H3 peptides.
- **SI Table 1B:** Averaged (AVG) and Standard Deviation (ST.DEV) of Relative Effective Concentration ($EC_{50}^{Rel}$) in Luminex assays to measure the interaction between GST-BPTF-PHD-BD constructs and PTM-defined histone H3 peptides.
- **SI Table 1C:** Relative Effective Concentration ($EC_{50}^{Rel}$) from Luminex assay between GST-tagged BPTF constructs and PTM-defined semi-synthetic nucleosomes.
- **SI Table 1D:** Averaged (AVG) and Standard Deviation (ST.DEV) of relative Effective Concentration ($EC_{50}^{Rel}$) from Luminex assay between GST-tagged BPTF constructs and PTM-defined semi-synthetic nucleosomes.
- **SI Table 2:** Settings used for intact nMS analysis of BPTF-bound nucleosome complexes (MS1).
- **SI Table 3:** Settings used for intact nMS analysis of CAPs (MS1).
- **SI Table 4:** Settings used for MS/MS fragmentation analysis of CAPs (MS2).
- **SI Table 5:** Settings used for characterization of ejected intact histone proteoforms from CAP-nucleosome complexes (MS1).
- **SI Table 6:** Settings used for MS/MS fragmentation analysis of histone proteoforms (MS2).
- **SI Table 7:** Settings used for Individual Ion Mass Spectrometry analysis of histone H3.2 proteoforms.

### Experimental Methods and Materials

**Protein expression and purification of recombinant chromatin reader constructs.** Human BPTF (Uniprot Q12830) PHD finger-Bromodomain (PHD-BD) cDNA was generated by PCR amplification and cloned into pGEX6p downstream of an N-terminal GST-tag and PreScission Protease cleavage site. Single amino acid substitutions were generated by Q5-site-directed mutagenesis (*NEB*) to produce BPTF mutants PHD* (W2891A), BD* (N3007A) and PHD*BD* (W2891A, N3007A).[1] Human DNMT3A (Uniprot Q9Y6K1) PWWP domain fused to Human MPP8 (Uniprot Q99549) Chromodomain, and *Populus trichocarpa* SHL (Uniprot Q9FEN9) Bromo-adjacent homology-Bromodomain (BAH-BD) cDNAs were generated by PCR amplification and cloned into pGEX-6P-2 downstream of an N-terminal GST-tag and PreScission Protease cleavage site. Human BRD4 (Uniprot O60855) BD1-BD2 cDNA was generated by PCR amplification and inserted by Gateway cloning (*Thermo Fisher Scientific*) to pDEST527 (*Addgene*: 11518) downstream of an N-terminal 6x-Histidine (6His) tag.

Recombinant BPTF, BRD4, DNMT3A-MPP8, and PtSHL constructs were transformed to *E. coli* (T7 Express *lysY*) (*NEB*). Liquid bacterial cultures were grown to $OD_{600}$ ~1, and protein expression induced using 0.8 mM IPTG at 18°C for 16 hours in LB media. Cells were pelleted by centrifugation and lysed in lysis buffer (25 mM Tris, pH 8.0, 200 mM NaCl, 10% Glycerol, 0.5% CHAPS, 1x EDTA-free Protease Inhibitor Cocktail (PIC), 2.5 mg/mL Lysozyme, 0.5 mM DTT, 1 µL of Universal Nuclease (25 kU; *Pierce*) / mL of cell lysate). Cell lysates were incubated with glutathione sepharose 4B (*Cytiva*), and beads washed three times with lysis buffer. Beads were incubated in glutathione elution buffer (10 mM reduced L-glutathione and 50 mM Tris HCl, pH 8.0) to purify recombinant GST-tagged BPTF, DNMT3A-MPP8, and PtSHL proteins. For purification of 6xHis-BRD4-BD1-BD2, cell lysates were mixed with Ni-NTA agarose (*Qiagen*) and beads washed three times using lysis buffer. Beads were incubated with lysis buffer supplemented with 300 mM Imidazole to purify 6xHis-BRD4. Purified chromatin-associated proteins were buffer exchanged to storage buffer (20 mM Tris pH 8.0, 200 mM NaCl, 20% Glycerol, 1 mM DTT) prior to aliquoting and freezing at -80ºC.

**Production and assembly of semi-synthetic nucleosomes.** Fully defined semi-synthetic mononucleosomes (*EpiCypher*) were produced as previously.[2,3] This study uses the recently proposed nucleosome nomenclature devised for accurate scientific communication in the chromatin and epigenetic fields.[4] Here ([H3K4me3]) indicates a fully defined semi-synthetic nucleosome where other positions not denoted are understood to be definitively unmodified. For native material, ({H3K4me3}) indicates a partially understood nucleoform where the named PTM is experimentally known to exist (as by immunoprecipitating H3K4me3 from a cell extract), but other sites of potential modification not denoted can be understood of undefined status.

**Cell Culture.** HeLa S3 cells (at confluence ~5-8x$10^5$ cells/mL) were cultured in DMEM supplemented with 1% Penicillin / Streptomycin and 10% HyClone Bovine Calf Serum (*Cytiva*) in spinner flasks in 5% CO2 incubator at 37°C.

**Extraction and purification of endogenous mononucleosomes.** HeLa S3 cell were collected and lysed under hypotonic condition with 2x pelleted cell volume (PCV) buffer A (10 mM HEPES, pH 7.9, 10 mM KCl, 1.5 mM $MgCl_2$, 340 mM sucrose, 10% Glycerol, 10 mM β-glycerophosphate, 5 mM Sodium butyrate, 0.5 mM DTT, and 1x EDTA-free PIC). Buffer A supplemented with 0.2% Triton X-100 was added in volume equal to cell suspension, and incubated on ice for 15 minutes. Cells were centrifuged at 1,300 x g for 5 minutes to release nuclei. Pelleted nuclei were washed with 2x PCV of buffer A and centrifuged at 1,300 x g for 15 minutes. Pelleted nuclei were resuspended in 2x PCV of buffer A. Nuclei suspension was supplemented with 2 mM CaCl and micrococcal nuclease (*NEB* M0247S) added (5 µL MNase / mL nuclei suspension). Suspension was incubated on a thermal mixer (*Eppendorf*) at 37°C for 30 minutes at 600 RPM. Digestion was quenched by addition of 2 mM EDTA and 1 mM EGTA and incubated on ice for 10 minutes. Nuclei suspension was supplemented with 0.05% Triton X-100 followed by 100 mM KCl and 300 mM NaCl in a dropwise fashion and incubated on ice for 30 minutes. Finally, nuclei suspension was clarified by centrifugation at 20,000 x g for 20 minutes at 4°C. The resulting supernatant containing soluble mononucleosomes were used for immunoprecipitation. DNA was purified from each mononucleosome extract (~10 µg) by Phenol-Chloroform extraction. Nucleosomal DNA (~150 bp) was resolved on a D1000 ScreenTape (*Agilent*) and analyzed on a TapeStation 4200 (*Agilent*).

**Purification of chromatin reader-bound endogenous nucleosome complexes.** Glutathione Sepharose 4B beads were saturated with excess GST-BPTF protein in 200 µL BPTF binding buffer (10 mM Tris-HCl pH 8.0, 300 mM KCl, 50ug/mL BSA, 1% NP-40, 1 mM DTT, 1x PIC), GST-PtSHL protein in PtSHL binding buffer (20 mM Tris pH 7.5, 150 mM NaCl, 0.01% Tween-20, 0.01% BSA), or GST-DNMT3A-MPP8 protein in DNMT3A-MPP8 binding buffer (20 mM Tris pH 7.5, 150 mM NaCl, 0.01% NP-40, 0.01% BSA + 1 mM DTT). Ni-NTA agarose beads (*Qiagen*) were saturated with excess 6xHis-BRD4 protein in BRD4 binding buffer (50 mM Tris-HCl pH 8.0, 100 mM NaCl, 0.1% NP-40). Beads were incubated at 4°C for two hours with rotation. Beads were washed four times with 500 µL of respective binding buffer. CAP-bound beads were incubated with 10 µg of endogenous mononucleosomes in 200 µL of respective binding buffer and incubated at either 4°C overnight or 25°C for two hours. Controls containing no CAPs were mixed with endogenous mononucleosomes to determine nonspecific binding to beads. Beads were washed three times with 500 µL of respective binding buffer to remove nonspecific interactions. CAP-endogenous nucleosome complexes were eluted under native conditions with either GST elution buffer (125 mM Tris-HCl pH 7.4, 50 mM reduced L-glutathione, 150 mM NaCl, 1 mM DTT, 1 mM EDTA) or His elution buffer (25 mM Tris pH 8.0, 100 mM NaCl, 200 mM Imidazole) on a thermal mixer at 37°C and 600 RPM for 30 minutes.

**Preparation of MagPlex-Nucleosome panels.** Luminex Avidin-conjugated MagPlex beads (16-distinct bead regions) were sourced (*Diasorin*) to assemble multiplexed nucleosome panels as previously.[5] All handling and incubations with MagPlex beads were performed under subdued lighting. Briefly, MagPlex beads were vortexed and incubated for 30 seconds in water bath sonicator. Beads were transferred to 1.5 mL tubes and pelleted using a magnet then washed twice with pre-conjugation buffer (50 mM Tris pH 7.5, 0.01% Tween-20). Nucleosome dilutions were prepared in pre-conjugation buffer. Thoroughly mixed beads were aliquot into 1.5 mL tubes and mixed with the assigned nucleosomes (*e.g.* MagPlex Region 13 & H3K4me3 Nuc) to 500 µL volume (5 μg Nuc / 1 million beads). Beads and nucleosomes were incubated on a rotator for 30 minutes at room temperature. Conjugated beads were then washed twice with post-conjugation buffer (50 mM Tris pH 7.5, 0.01% Tween-20, 0.01% BSA) and resuspended to ~1.5 M beads/mL. Bead concentrations were determined using a Luna cell counter (*Logos Biosystems*); >90% monodispersed beads). Bead concentrations were adjusted to 1M/mL using post-conjugation buffer. A master mix was then assembled with equal quantity (*e.g.*, 500K beads) of each bead region. The master mix was pelleted on a magnet, decanted, and resuspended at 1M/mL per region in storage buffer (10 mM cacodylate pH 7.5, 0.01% BSA, 0.01% Tween-20, 1 mM EDTA, 10 mM beta mercaptoethanol, 50% glycerol) and stored at -20°C. For quality control, each nucleosome PTM was detected with previously characterized anti-PTM antibodies. Briefly, 50 μL of the multiplex bead panel (20,000 beads/mL/region; 1000 beads/region) was combined with 50 μL of antibody (1:125, 1:500, 1:2000) in a 96-well plate and incubated 60 minutes with shaking to maintain bead resuspension. Bead and antibody dilutions were prepared in QC assay buffer (50 mM Tris pH 7.5, 250 mM NaCl, 0.01% Tween-20, 0.01% BSA). Beads were then washed for three cycles on a magnet using 100 μL of QC assay buffer and shaken for 2 minutes between each cycle. Anti-IgG PE [Anti-Rabbit (*Biolegend* 406421) or Anti-Mouse (*Biolegend* 405307)] were diluted 1:100 in QC assay buffer and added to each well and followed by incubation for 30 minutes with shaking. Beads were then washed for two cycles on a magnet using 100 μL of QC assay buffer and shaken for 2 minutes between each cycle. Lastly, beads were resuspended in 100 μL QC assay buffer and fluorescence measured using the FlexMap3D Instrument (*Diasorin*).

**Luminex-based dCypher binding assays.** dCypher (Luminex) binding assays were performed by combining 50 μL of multiplexed beads (20,000 beads/mL/region; 1000 beads/region) with 50 μL of tagged query domain (at concentrations noted in figures) in a 96-well plate.[3,5] Query protein and beads were diluted in assay buffer (20 mM Tris pH 7.5, 250 mM NaCl, 0.01% BSA, 0.01% Tween-20, 1 mM DTT). The reaction plate was incubated for 60 minutes with shaking to maintain bead resuspension. Beads were washed for three cycles on a magnet using 100 μL of assay buffer and shaken for 2 minutes between each cycle. 100 μL of Anti-Tag antibody [1:2000 Anti-GST (*Fortis* A190-122A)] was added to each well and incubated for 30 minutes while shaking. Beads were then washed for three cycles on a magnet using 100 µL of assay buffer and shaken for 2 minutes between each cycle. Anti-IgG PE [Anti-Rabbit (*Biolegend* 406421)] was diluted 1:100 in assay buffer and added to each well and incubated 30 minutes with shaking. Beads were then washed for two cycles on a magnet using 100 µL of assay buffer and shaken for 2 minutes between each cycle. Lastly, beads were resuspended in 100 µL assay buffer and fluorescence measured using the FlexMap3D Instrument (*Diasorin*). Single concentration bar graphs or binding curves generated using a non-linear 4PL curve fit were plotted in GraphPad Prism 10 (v10.10.0).

**Immunoblotting.** Immunoblotting was performed to determine pulldowns of reader-endogenous nucleosome complexes. The following primary antibodies were used: Anti-GST tag (*Invitrogen* MA4-004, 1:1000), Anti-6xHis tag (*Invitrogen* 37-2900), and Anti-Histone H3 (*Cell Signaling Technologies* 4499S, 1:2000). Secondary antibodies used were: Anti-Mouse-HRP (*Jackson ImmunoResearch* 115-035-044, 1:2000) and Anti-Rabbit-HRP (*Jackson ImmunoResearch* 111-035-003, 1:2000).

**Native Mass Spectrometry.** Recombinant BPTF proteins were incubated with PTM-defined semi synthetic nucleosomes in equimolar concentrations (1 µM) in binding buffer (20 mM HEPES pH 7.5, 250 mM NaCl, 0.01% BSA, 0.01% NP-40, 1 mM DTT) for 1 hour at 4°C. The mixture was subsequently desalted, and buffer exchanged using 30-kDa molecular-weight-cut-off (MWCO) spin filters (*MilliporeSigma* UFC503096) into 150 mM Ammonium acetate (AmAc) prepared in LC/MS grade Optima water (*Thermo Fisher Scientific* AAB-W6-4) as previously.[6] Briefly, spin filters were first equilibrated with 500 µL water, and spun for 3 minutes at 10,000 x *g*. Samples were loaded into equilibrated spin filter and centrifuged at 10,000 x *g* for 10 minutes at 4°C. The buffer exchange process was repeated up to ten times to allow sufficient removal of MS-incompatible components. In each stage, the solution was spun down to <100 µL, and the filter replenished up to 500 µL with 150 mM AmAc. Buffer exchange of reader-endogenous nucleosome complexes into 150 mM AmAc were performed under the same procedure. Samples were concentrated to 1 µM prior to MS analysis. Samples were loaded into static NSI tips (*Thermo Fisher Scientific* ES380) for native electrospray ionization using a

NanoSpray Flex Ion Source (*Thermo Fisher Scientific*). For PTM-defined semi-synthetic nucleosome and BPTF binding experiments, Nuc-MS analysis was performed on an Orbitrap Q Exactive Ultra High Mass Range (UHMR) MS (*Thermo Fisher Scientific*).[7] For analysis of chromatin-associated proteins (CAPs) or CAP-endogenous nuc complexes, Nuc-MS analysis was performed on an Orbitrap Ascend Tribrid MS (*Thermo Fisher Scientific*). In-depth details pertaining to instrumentation and MS/MS parameters can be found in **SI Table 2-6**.

**Single Ion Data Acquisition and Analysis for Individual Ion Mass Spectrometry (I2MS).** I2MS[8–10] allows for the direct charge state assignment of individual ions using an Orbitrap-based mass analyzer containing a harmonic potential. Here it was performed on an Orbitrap Eclipse Tribrid (*Thermo Fisher Scientific*), and used to characterize endogenous histone H3.2 from ejected CAP-endogenous nucleosome complexes and I2MS In brief, histone subunits were first released from intact complexes using in-source collision-induced-dissociation (CID) at 250 eV. Histone H3.2 proteoform precursor ions were subsequently fragmented via electron transfer dissociation (ETD). The dissociation condition was optimized to retain ~ 50% of precursor ions. Injection times were manually adjusted to resolve individual fragment ions. Charge assignment for individual ions was performed using $I^2$MS Processing Pipeline with an ion lifetime threshold of 210 ms, a signal-to-noise threshold of 3, a minimum bin size of 1, and a voting probability threshold of 0.5. Fragment ion annotation was performed with a TDValidator (*Proteinaceous*, v1.1.242332.1) with manual confirmation. In-depth details pertaining to instrumentation and MS/MS parameters can be found in **SI Table 7**.

**Mass Spectrometry data analysis.** FreeStyle (*Thermo Fisher*, v1.8) was used to obtain time-dependent acquisition of MS spectra. UniDec (v 6.0.4)[11] was used to generate deconvoluted MS1 spectra. For data processing the following settings were used: *m/z* 4000-12000. For UniDec parameters the following settings were used: Charge range: 1-50+, Mass range: 5000-350,000 kDa, Sample mass every 10 Da, Smooth Nearby Points: Some, Suppress Artifacts: None, Peak FWHM: 0.85, Charge Smooth width: 1.0, Point Smooth Width: 1.0, and Maximum # of iterations: 100. TDValidator (v1.1.242332.1) was used to characterize intact masses of histone subunits at isotopically resolution (MS1) and fragmentation data of histones and reader proteins (MS2).[12] The following TDValidator Precursor parameters for intact mass analysis were as follows: Max PPM Tolerance: 20.00, Sub PPM Tolerance: 20.00, Cluster Tolerance: 0.70, Charge Range: 5-50, Minimum Score: 0.30, signal-to-noise cutoff: 3.00, Mercury7 Limit: 0.0001, and Isotopically Resolved. Relative quantification of intact histone proteoforms were performed by extracting signal intensities (MS1) at intact precursor level and comparisons of histone proteoform between bulk HeLa and reader-enriched nucleosomes were performed using the same charge states (z). The following TDValidator Fragments parameters for high resolution fragmentation data were as follows: Max PPM tolerance: 10-15 ppm, Sub PPM tolerance: 5 ppm, Charge Range: 1-15, Minimum Score: 0.35-0.70, signal-to-noise cutoff: 3.00, Minimum Size: 1, Mercury7 Limit: 0.0001, and FDR Decoy Runs: 1000. ProSight Lite (v1.4) and TDValidator were used to match *b* and *y*, or *c* and *z* fragment ions corresponding to histone or reader sequences and generate P Scores for each histone proteoform. All mass spectrometry .raw files were uploaded to MassIVE (MSV000097336).

**Safety Statement:** No unexpected or unusually high safety hazards were encountered.

(4) Keogh, M.-C.; Almouzni, G.; Andrews, A. J.; Armache, K.-J.; Arrowsmith, C. H.; Baek, S. H.; Bedford, M. T.; Bernstein, E.; David, Y.; Denu, J. M.; Garcia, B. A.; Glass, K. C.; Helin, K.; Henikoff, S.; Jensen, O. N.; Kelleher, N. L.; Kutateladze, T. G.; Lindner, H.; Lu, C.; Josefowitz, S. Z.; Luger, K.; Mallick, P.; Musselman, C. A.; Muir, T. W.; Paša-Tolić, L.; Schneider, R.; Shi, X.; Sidoli, S.; Tyler, J. K.; Wolberger, C.; Workman, J. L.; Strahl, B. D.; Young, N. L.; Young@bcm, N. *A Needed Nomenclature for Nucleosomes*.

(12) Fornelli, L.; Srzentić, K.; Huguet, R.; Mullen, C.; Sharma, S.; Zabrouskov, V.; Fellers, R. T.; Durbin, K. R.; Compton, P. D.; Kelleher, N. L. Accurate Sequence Analysis of a Monoclonal Antibody by Top-Down

and Middle-Down Orbitrap Mass Spectrometry Applying Multiple Ion Activation Techniques. *Anal Chem* **2018**, *90* (14), 8421–8429. https://doi.org/10.1021/acs.analchem.8b00984.

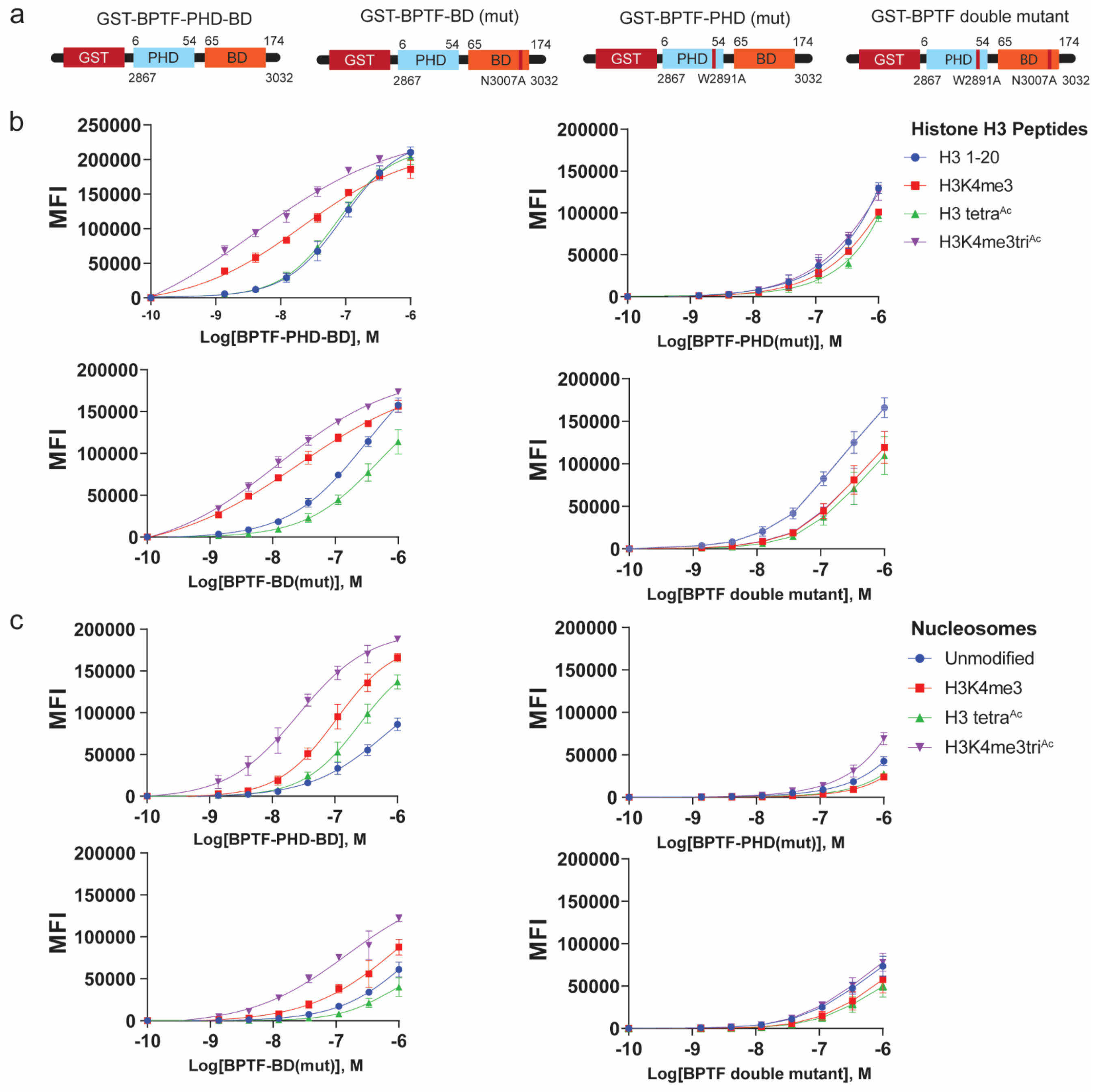


**SI Figure 1: Differential BPTF PHD-BD binding preference for histone peptides and nucleosomes. a)** N-terminally GST-tagged BPTF (Uniprot Q12830) constructs (including domain coverage and mutant position). **b-c)** Binding (by dCypher Luminex assay) of each BPTF-PHD-BD (titrated Queries) to PTM-defined histone peptides **(b)** or semi-synthetic nucleosomes **(c)** (each fixed concentration Targets). Data is presented as Measured Median Fluorescence Intensity (MFI) as a function of logarithmic transformation of BPTF molar concentration (M). Experiments were performed with three independent biological replicates. Error bars represent standard deviation from mean.

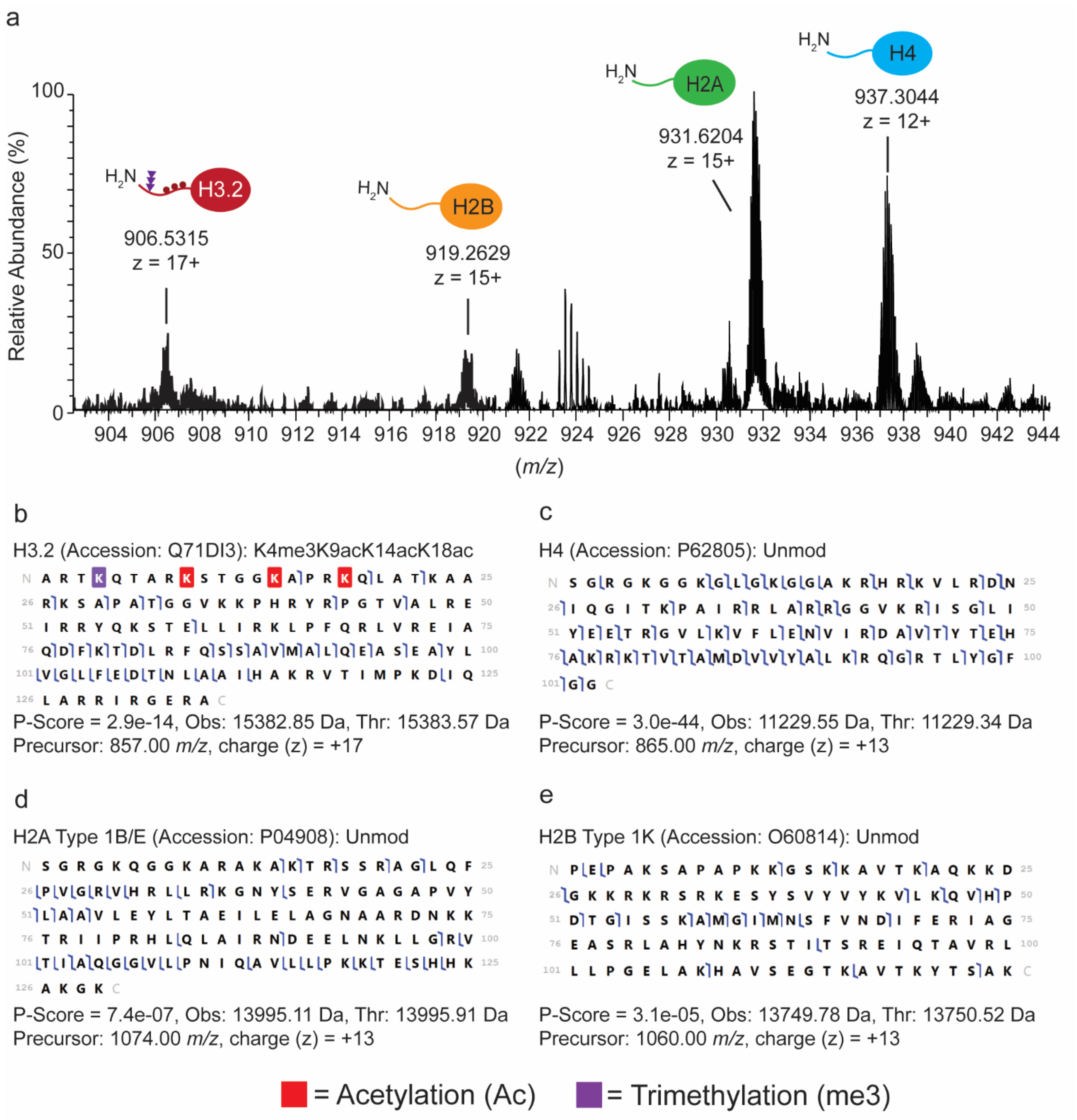


**SI Figure 2: Nuc-MS characterization of BPTF-bound ([H3K4me3tri$^{Ac}$]) nucleosomes. a)** Representative MS1 spectra of intact histones ejected from GST-BPTF-PHD-BD-bound ([H3K4me3K9acK14acK18ac]) nucleosomes. **b-e)** Graphical fragment maps of the most abundant intact histone precursor ions: H3K4me3K9acK14acK18ac **(b)**, H4 **(c)**, H2A Type 1B/E **(d)**, and H2B Type 1K **(e)**. Forward and reverse blue flags represent *b* and *y* ions resulting from higher-energy collisional dissociation (HCD), respectively. Experiments were conducted with three independent biological replicates. Fragments were manually validated using TDValidator and corresponding P-Scores were calculated using ProSight Lite.

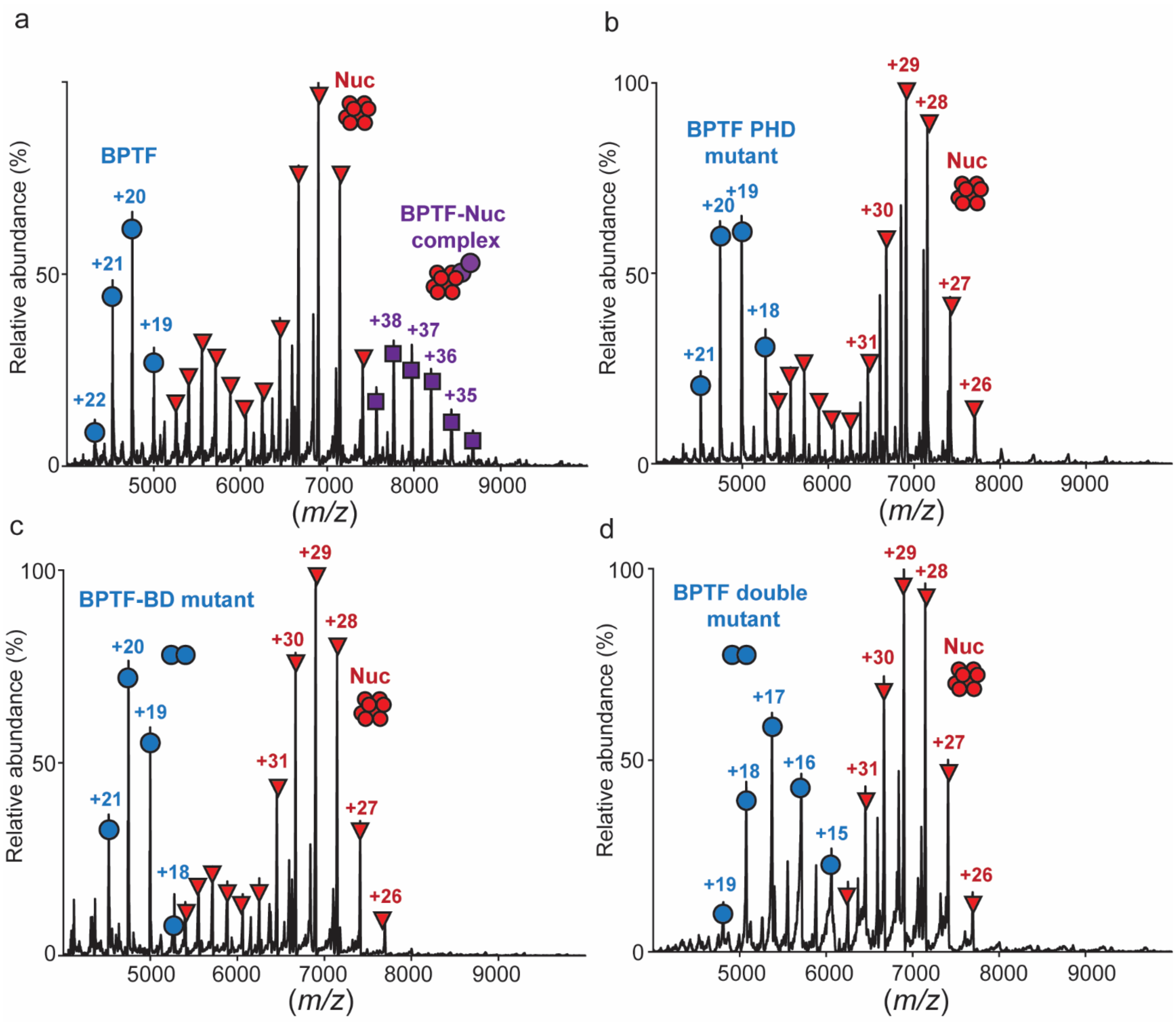


**SI Figure 3: Both domains in the BPTF-PHD-BD tandem are required for effective binding to ([H3K4me3tri$^{Ac}$]) nucleosomes. a-d)** Representative Native MS1 spectra of ([H3K4me3K9acK14acK18ac]) mononucleosomes mixed with either BPTF PHD-BD **(a)** or the loss-of-function mutants PHD* **(b)**, BD* **(c)** or PHD*BD* **(d)**. Experiments were conducted with three independent biological replicates.

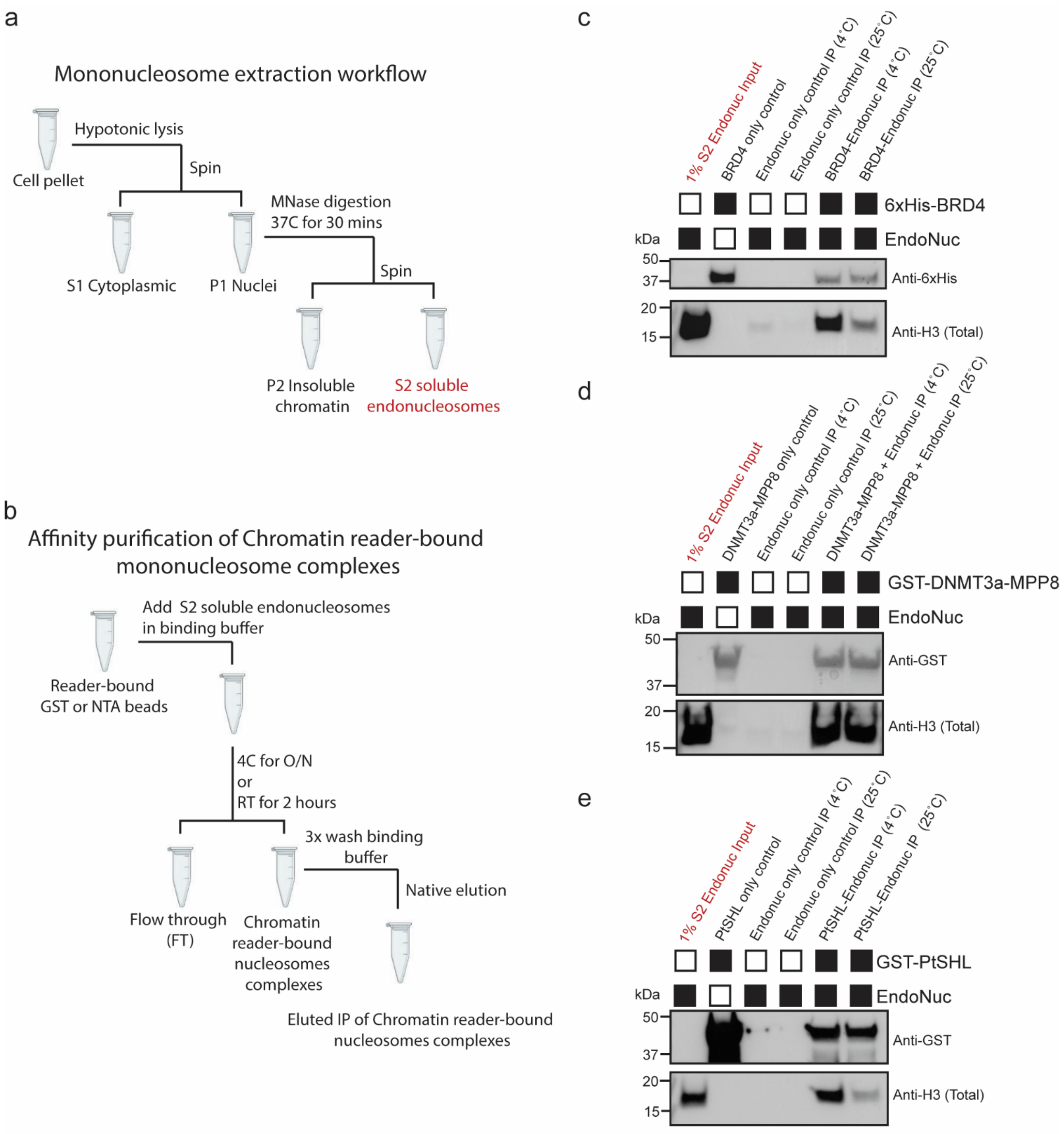


**SI Figure 4: Extraction and affinity purification of CAPs complexed with endogenous nucleosomes. a)** Graphical representation of mononucleosome extraction from HeLa cells. **b)** Graphical representation of workflow to enrich for CAP:nucleosome (CAP:nuc) complexes. **c-e)** Representative affinity purification of tandem reader bound endogenous nucleosomes: via 6xHis-BRD4-BD1-BD2 **(c)**, GST-DNMT3A-PWWP-MPP8-PHD **(d)**, or GST-PtSHL-BAH-PHD **(e)**. All affinity purifications were performed with three independent biological replicates.

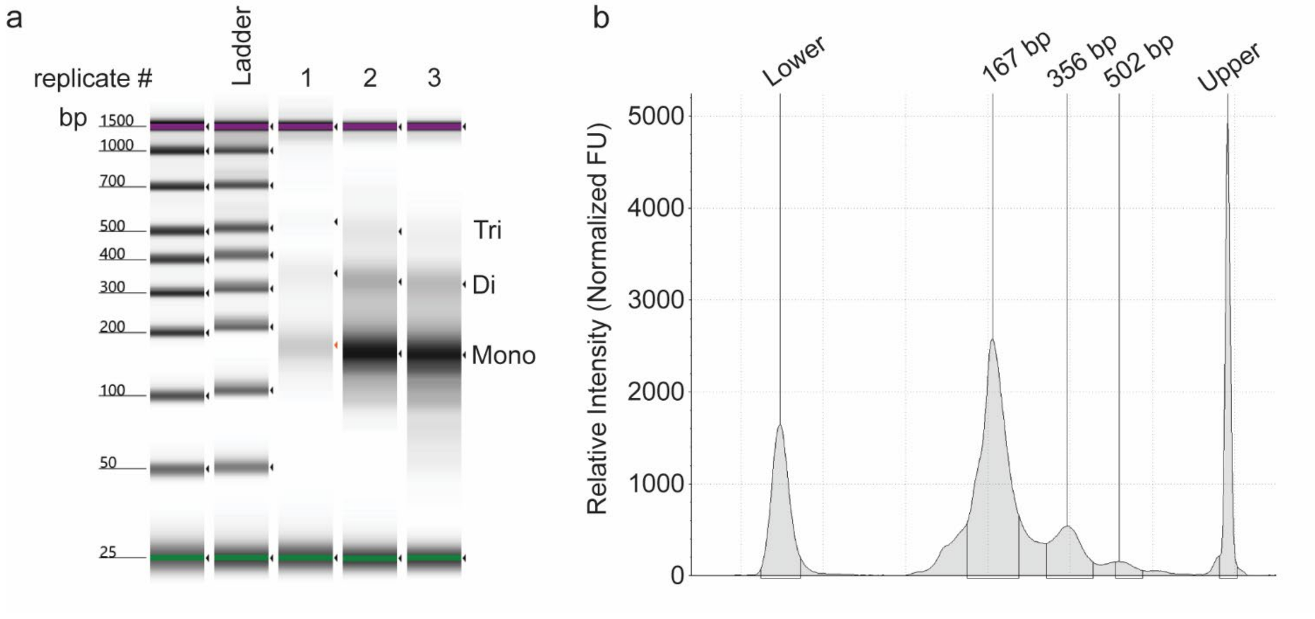


**SI Figure 5: Purification of nucleosomal DNA from endogenous HeLa mononucleosomes. a)** Electrophoretic gel separation of purified nucleosomal DNA fragments originated from endogenous nucleosomes (mono-, di- and tri-) extracted from HeLa cells. **b)** Relative abundance of DNA fragments as Normalized Fluoresence Units (FU).

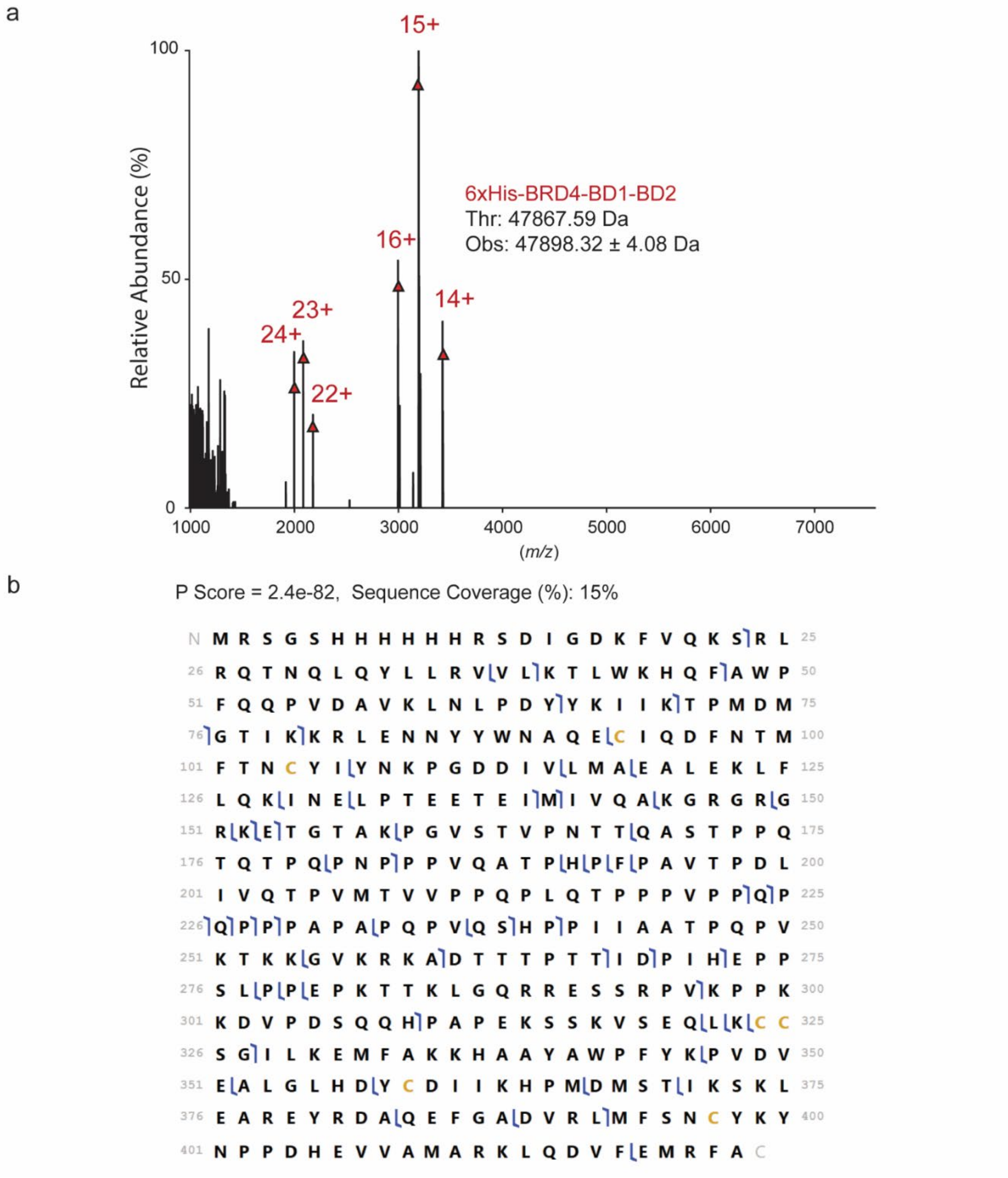


**SI Figure 6: Characterization of 6xHis-BRD4-BD1-BD2 native tandem reader.** **a)** Representative Native MS1 analysis of 6xHis-BRD4-BD1-BD2. Charge state distributions of intact 6xHis-BRD4-BD1-BD2 are represented in red (+14 to +16) and (+22 to +24). **b)** Representative graphical fragment map of isolated 6xHis-BRD4-BD1-BD2 (precursor ion: 3141.00 *m/z*) fragmented by higher-energy collisional dissociation (HCD). Forward and reverse blue flags respectively represent *b* and *y* ions. Experiments were conducted with three independent biological replicates. Fragments were manually validated using TDValidator and corresponding P-scores were calculated using ProSight Lite.

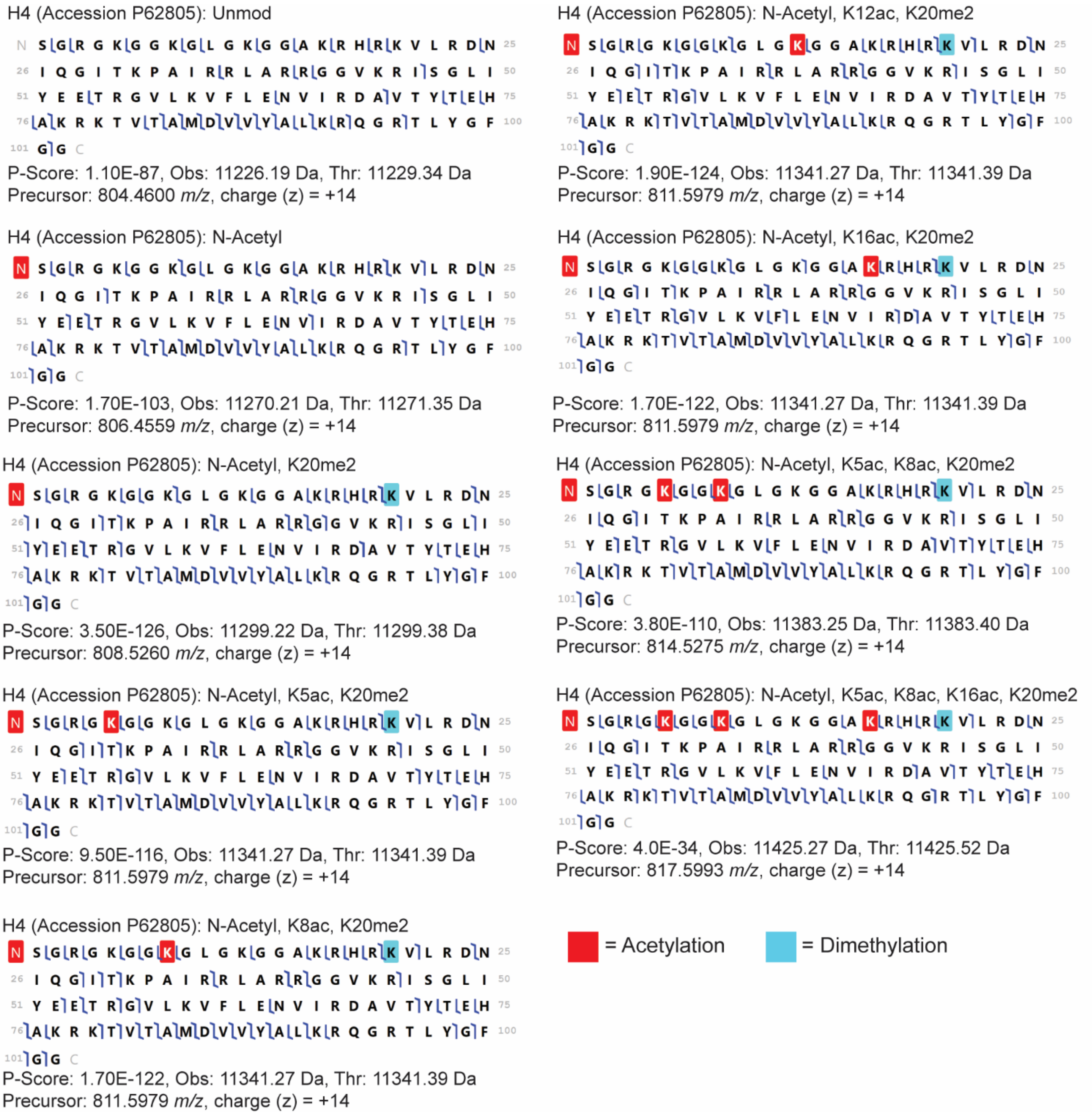


**SI Figure 7: Characterization of Histone H4 proteoforms from BRD4-enriched endogenous nucleosomes.** Representative graphical fragment maps of tandem MS fragmentation of histone H4 proteoforms in BRD4-enriched nucleosomes. Tandem MS fragmentation was performed using higher-collisional energy dissociation (HCD) across a distribution of H4 proteoforms (precursor ions (*m/z*) at charge state (z) +14 represent the intact H4 proteoforms used for tandem MS). Forward and reverse blue flags respectively represent *b* and *y* ions. Measurements were performed with three independent biological replicates. Observed (Obs) and theoretical (Thr) are represented as monoisotopic masses (Da). Fragments were manually validated using TDValidator and corresponding P-scores were calculated using ProSight Lite.

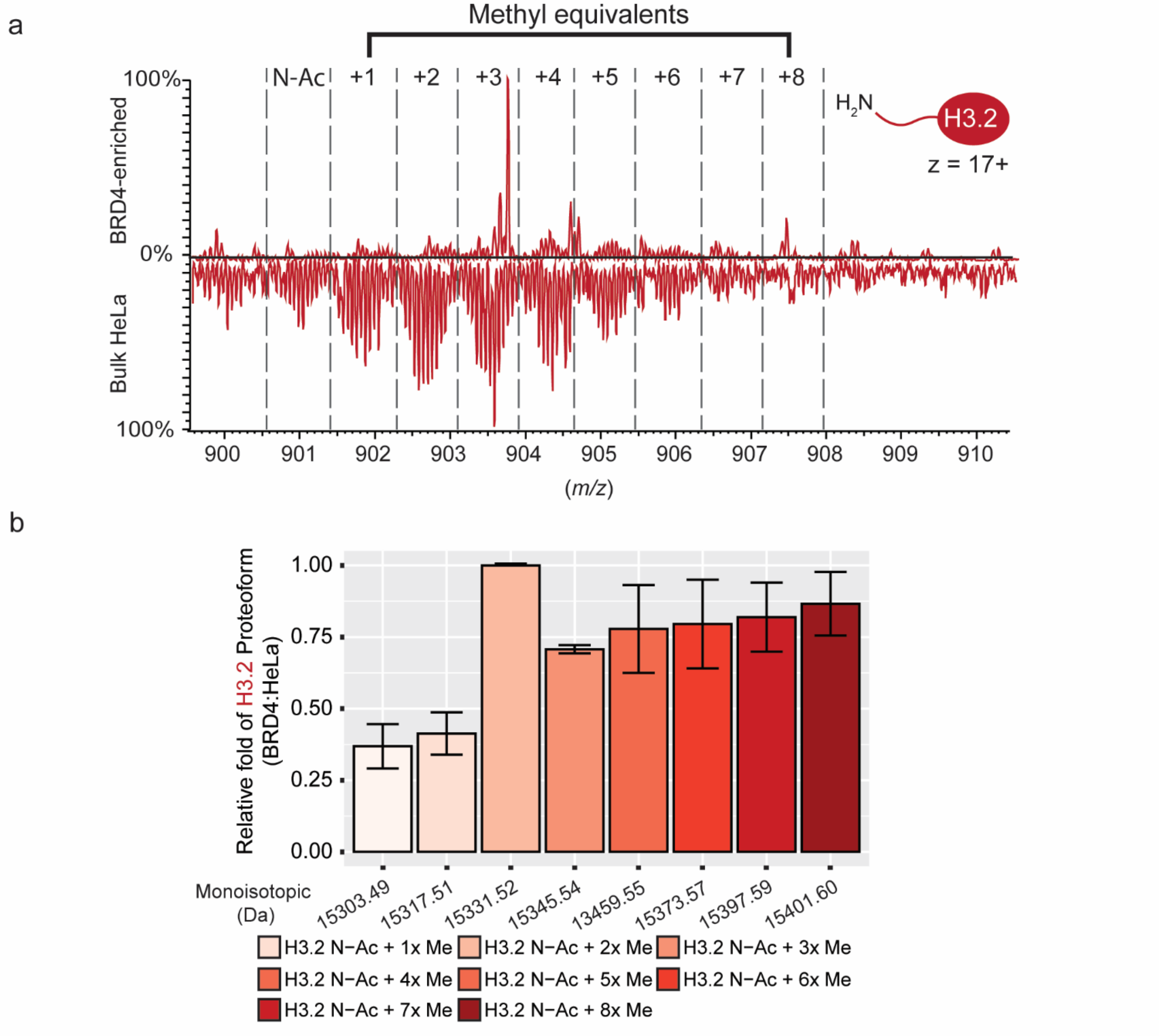


**SI Figure 8: Relative quantification of H3.2 proteoforms from BRD4-enriched endogenous nucleosomes. a-b)** Representative MS1 spectra **(a)** and quantification **(b)** of histone H3.2 proteoforms. Methyl equivalents are represented as (nxMe). Quantification of enrichment was calculated by determining the relative abundances of each histone proteoform and normalizing to H3.2: N-Ac + 3xMe. The relative ratio of BRD4:HeLa bulk is the normalized values of histone proteoform enriched by BRD4 divided by histone H4 proteoform in HeLa bulk. Error bars represent standard deviation from the mean.

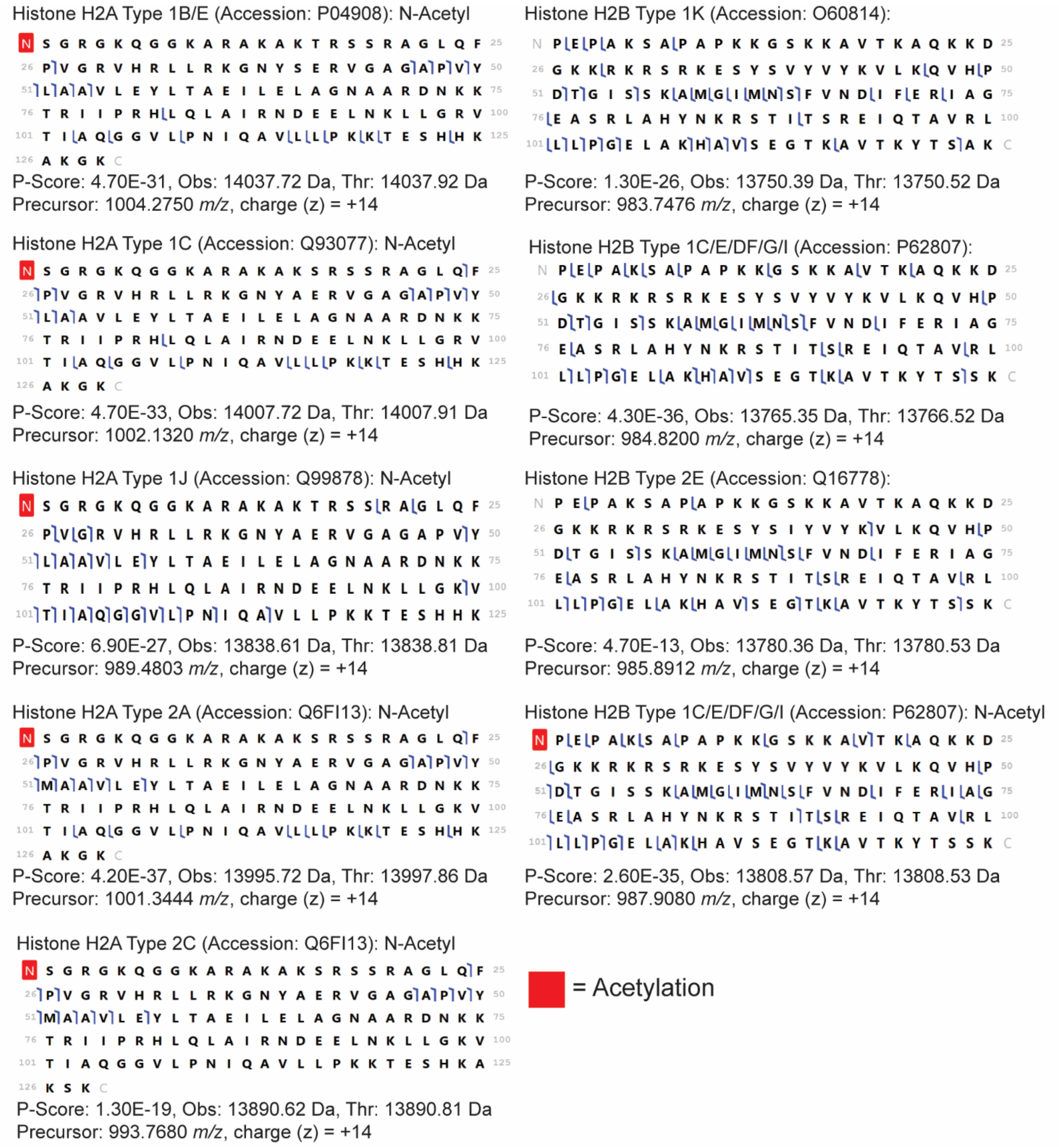


**SI Figure 9: Characterization of Histone H2A and H2B proteoforms from BRD4-enriched endogenous nucleosomes.** Representative graphical fragment maps of tandem MS fragmentation of histone H2A and H2B proteoforms in BRD4-enriched nucleosomes. Tandem MS fragmentation was performed using higher-collisional energy dissociation (HCD) across a distribution of H2A and H2B proteoforms (precursor ions (*m/z*) at charge state (z) +14 represent the intact H2A and H2B proteoforms used for tandem MS). Forward and reverse blue flags respectively represent *b* and *y* ions. Measurements were performed with three independent biological replicates. Observed (Obs) and theoretical (Thr) are represented as monoisotopic masses (Da). Fragments were manually validated using TDValidator and corresponding P-scores were calculated using ProSight Lite.

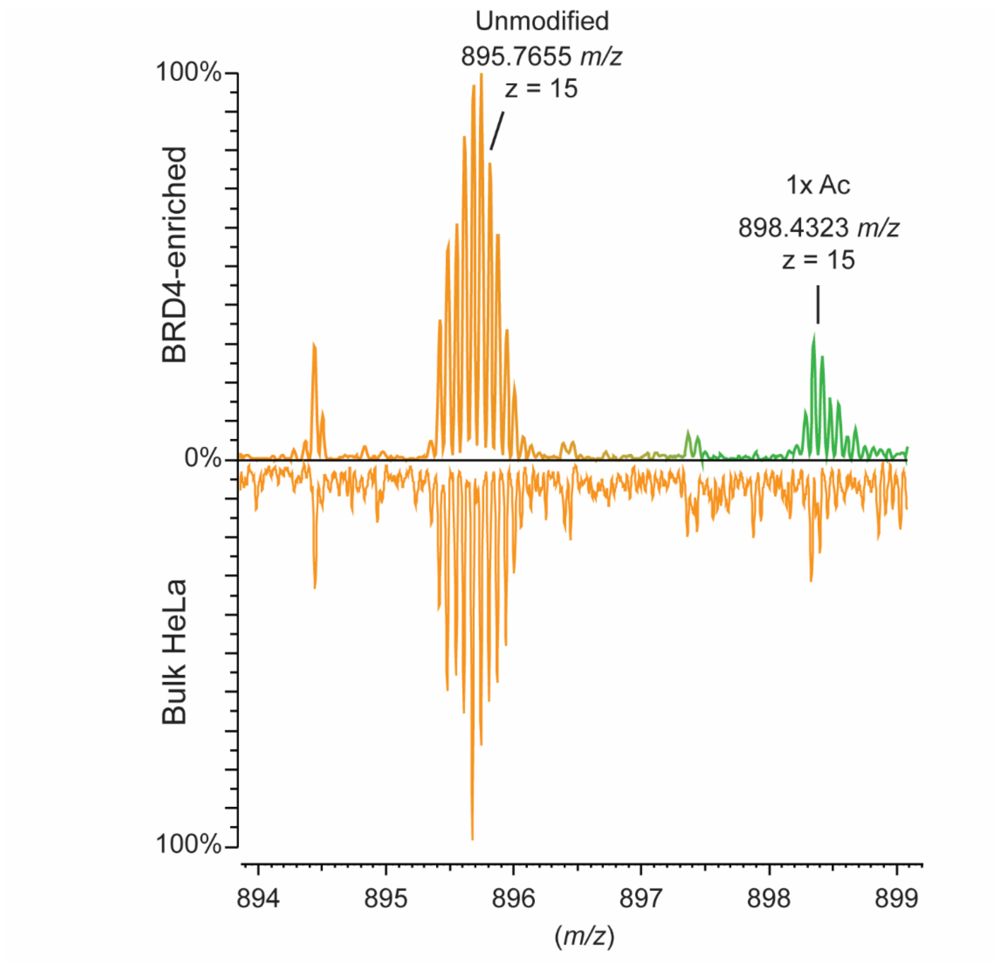


**SI Figure 10: BRD4 enriches for acetylated histone H2A.Z variant.** Representative MS1 spectra of intact histone H2A.Z variant proteoform landscape ejected from nucleosomes enriched by BRD4 (top) or from bulk HeLa endogenous nucleosomes (bottom). Highlighted in green is an acetylated H2A.Z. Experiments were conducted with three independent biological replicates.

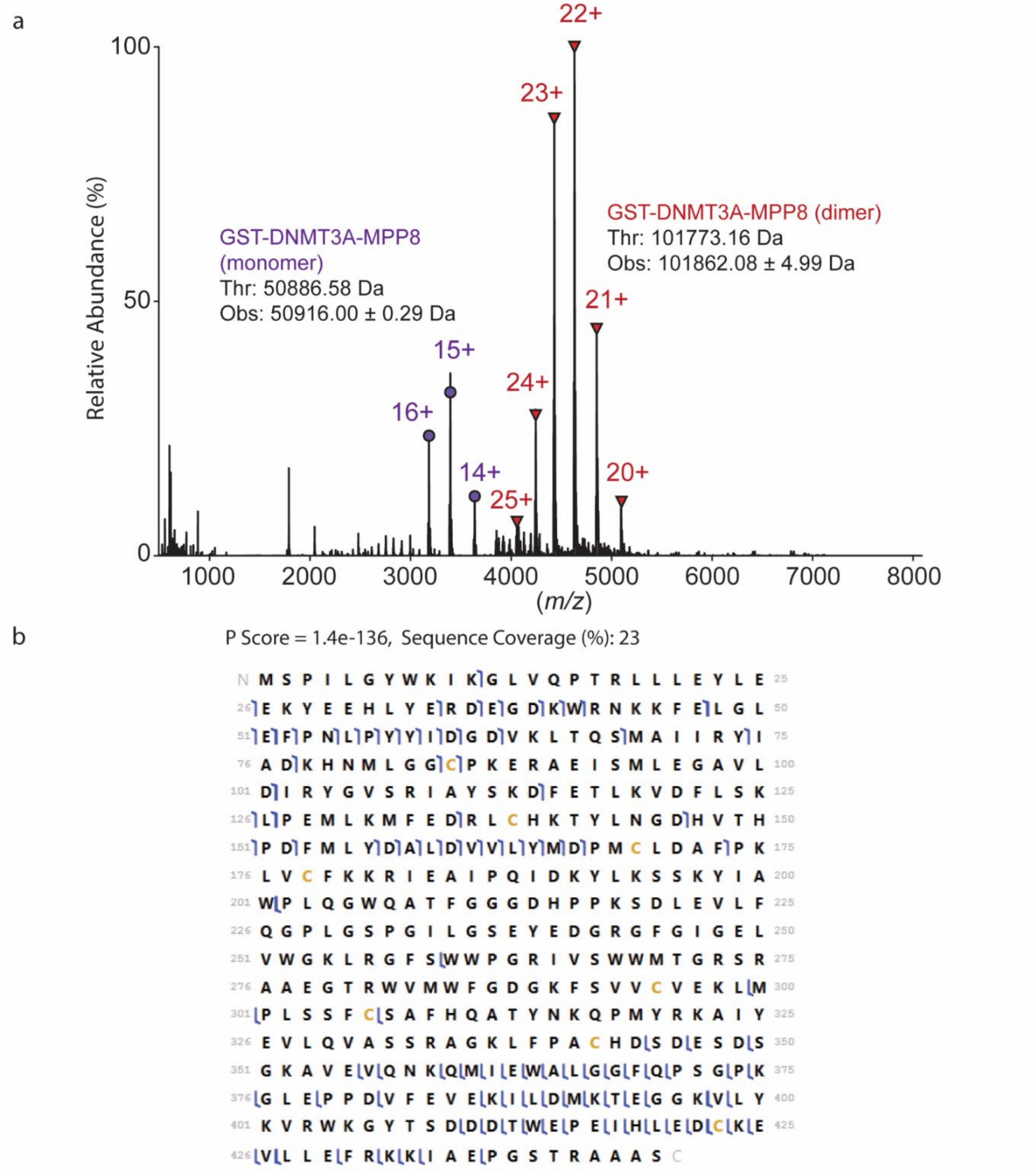


**SI Figure 11: Characterization of GST-DNMT3A-MPP8 PWWP-CD chimeric tandem reader. a)** Representative Native MS1 analysis of GST-DNMT3A-MPP8 PWWP-Chromodomain chimera tandem reader dimer (possibly via GST-tag) and monomer showing respective charge state distributions in red (+20 to +25) or purple (+14 to +16), respectively. **b)** Representative graphical fragment map of isolated GST-DNMT3A-MPP8 PWWP-CD (precursor ion: 4630.00 *m/z*) fragmented by higher-energy collisional dissociation (HCD). Forward and reverse blue flags respectively represent *b* and *y* ions. Experiments were conducted with three independent biological replicates. Fragments were manually validated using TDValidator and corresponding P-scores were calculated using ProSight Lite.

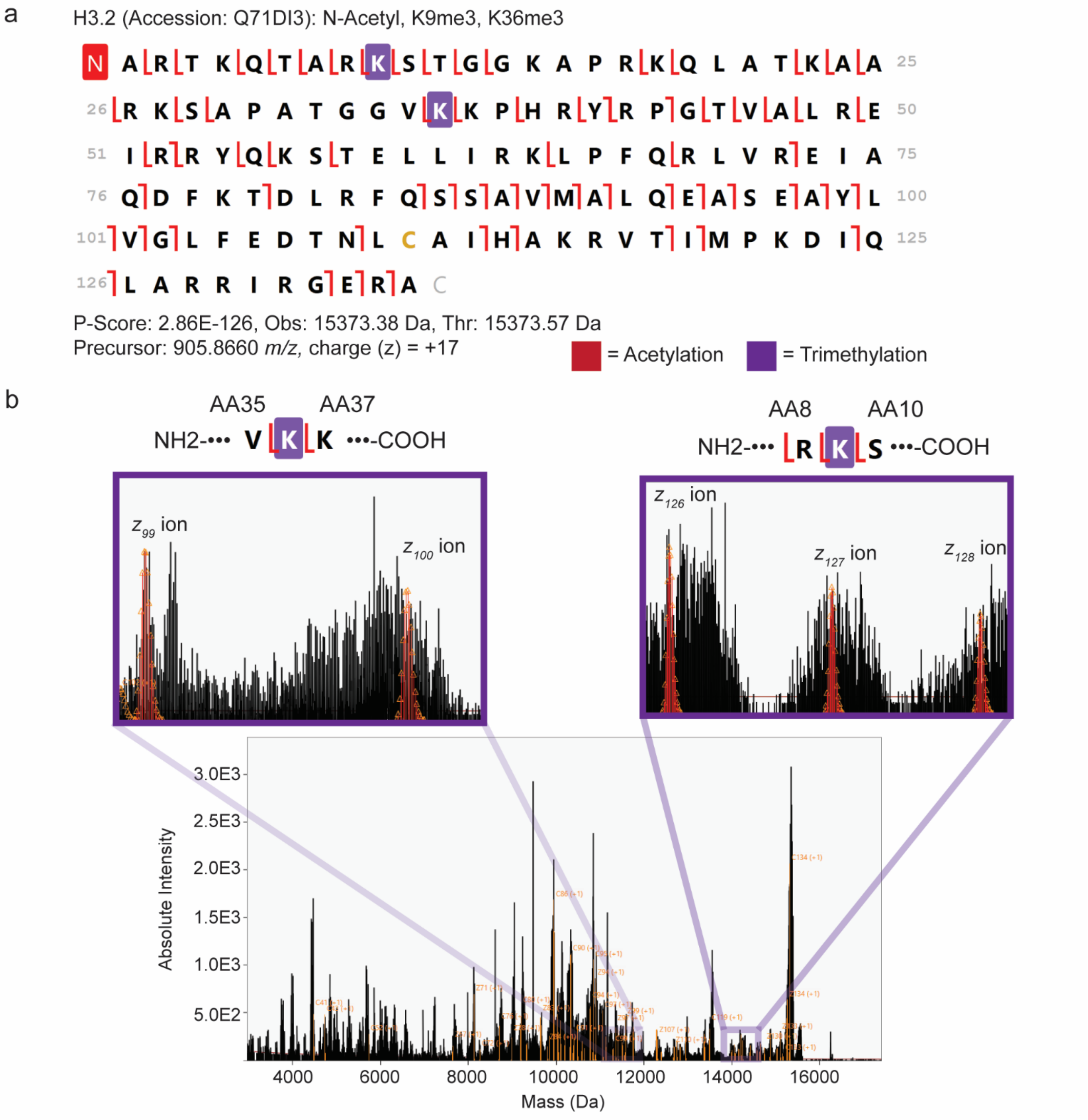


**SI Figure 12: Characterization of {H3.2 N-Acetyl, K9me3, K36me3} proteoform from DNMT3A-MPP8-enriched endogenous nucleosomes. a)** Representative graphical fragment map of tandem MS fragmentation of {H3.2 N-Acetyl, K9me3, K36me3} proteoform. Tandem MS fragmentation was performed using electron-transfer dissociation (ETD). Forward and reverse red flags respectively represent *c* and *z* ions. **b)** Deconvoluted mass spectra (Da) of tandem MS fragments of {H3.2 N-Acetyl, K9me3, K36me3} proteoform. Orange represents matched fragment ions. Insets represent fragment ions localizing to K9me3 and K36me3. Measurements were performed at three independent biological replicates. Observed (Obs) and theoretical (Thr) are represented as monoisotopic masses (Da). Fragments were manually validated using TDValidator and corresponding P-scores were calculated using ProSight Lite.

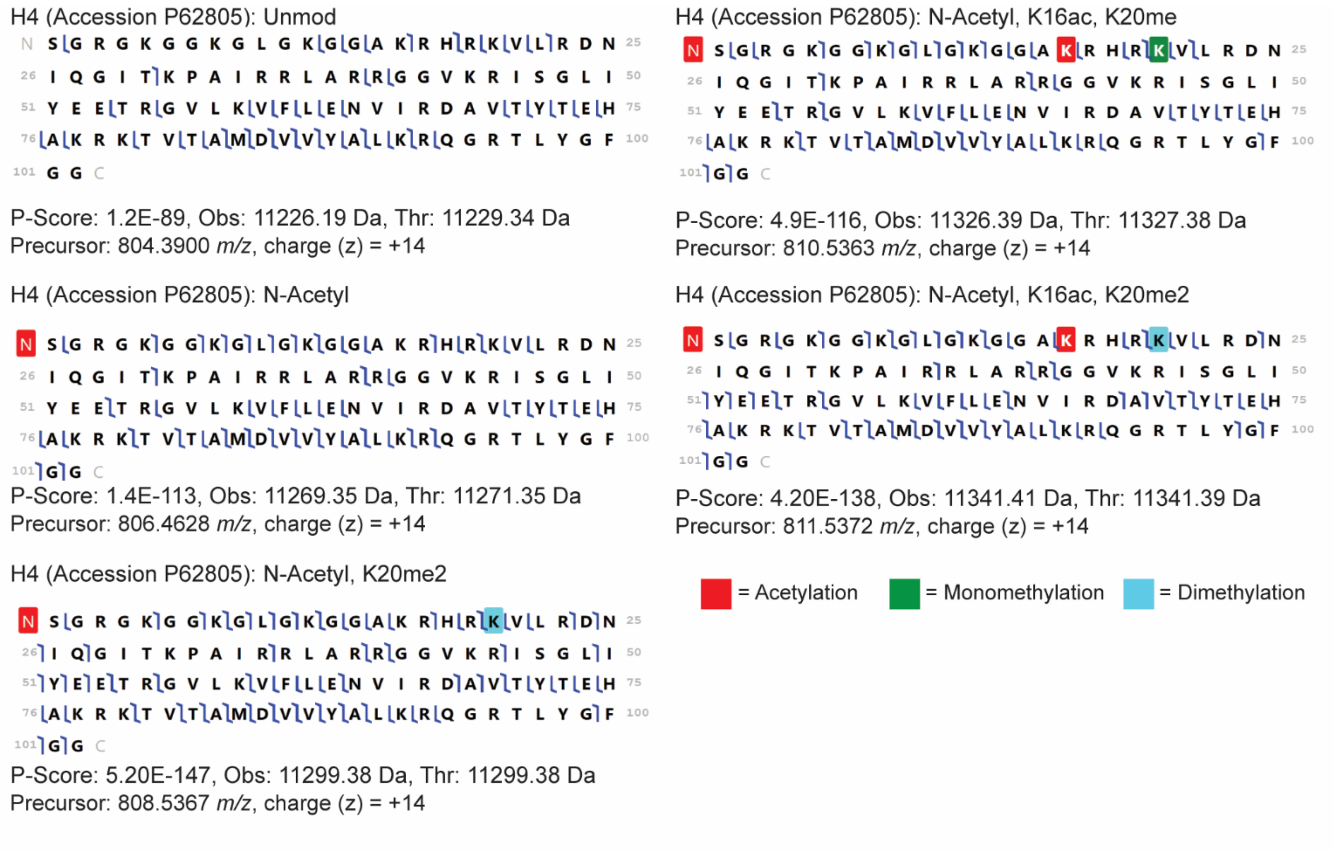


**SI Figure 13: Characterization of Histone H4 proteoforms from DNMT3A-MPP8-enriched endogenous nucleosomes by MS/MS fragmentation.** Representative graphical fragment maps of tandem MS fragmentation of histone H4 proteoforms in DNMT3A-MPP8-enriched nucleosomes. Tandem MS fragmentation was performed using higher-collisional energy dissociation (HCD) across a distribution of H4 proteoforms (precursor ions (*m/z*) at charge state (z) +14 represent the intact H4 proteoforms used for tandem MS). Forward and reverse blue flags respectively represent *b* and *y* ions. Measurements were performed at three independent biological replicates. Observed (Obs) and theoretical (Thr) are represented as monoisotopic masses (Da). Fragments were manually validated using TDValidator and corresponding P-scores were calculated using ProSight Lite.

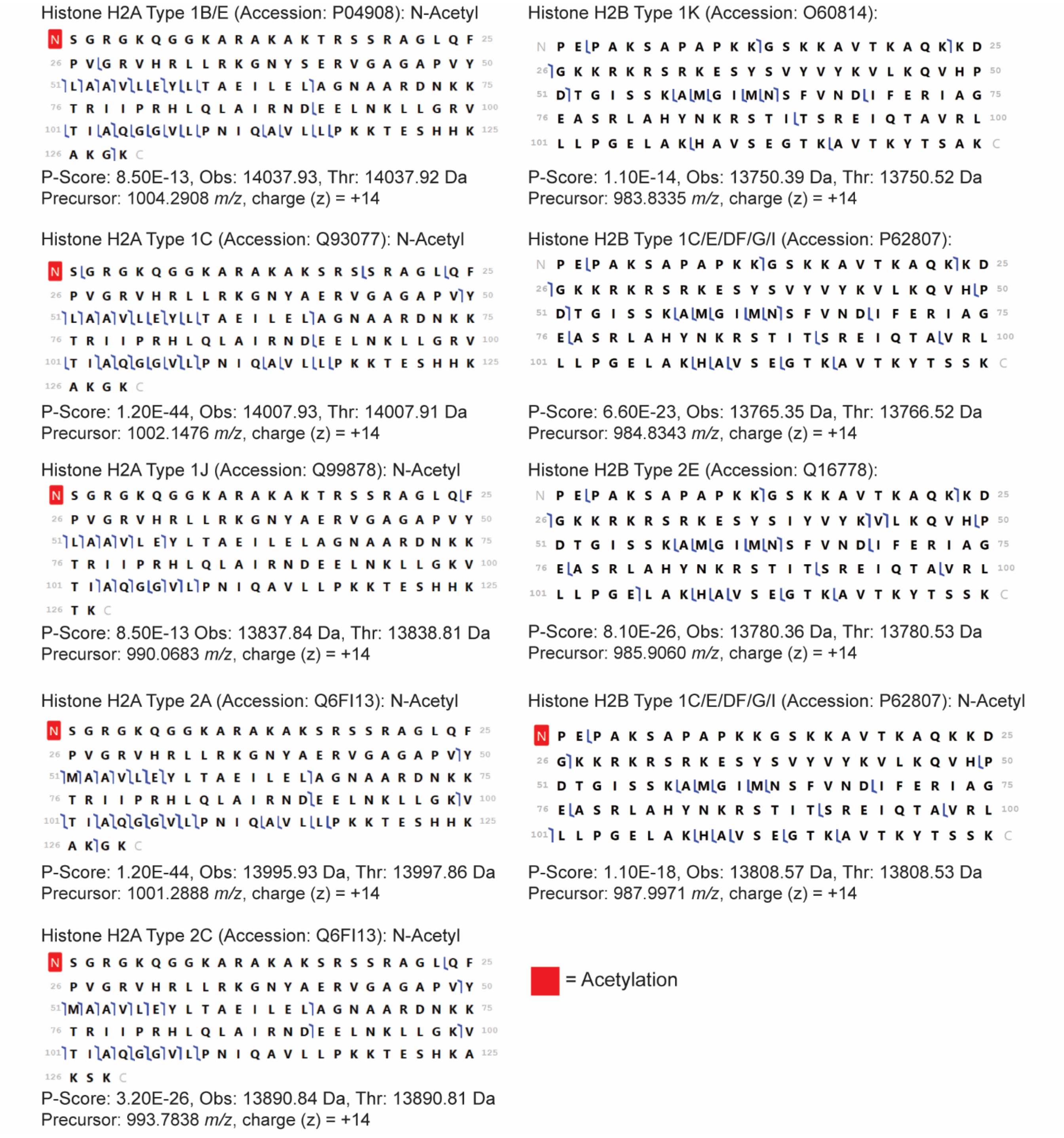


**SI Figure 14: Characterization of Histone H2A and H2B proteoforms from DNMT3A-MPP8-enriched endogenous nucleosomes.** Representative graphical fragment maps of tandem MS fragmentation of histone H2A and H2B proteoforms in DNMT3A-MPP8-enriched nucleosomes. Tandem MS fragmentation was performed using higher-collisional energy dissociation (HCD) across a distribution of H2A and H2B proteoforms (precursor ions (*m/z*) at charge state (z) +14 represent the intact H2A and H2B proteoforms used for tandem MS). Forward and reverse blue flags respectively represent *b* and *y* ions. Measurements were performed with three independent biological replicates. Observed (Obs) and theoretical (Thr) are represented as monoisotopic masses (Da). Fragments were manually validated using TDValidator and corresponding P-scores were calculated using ProSight Lite.

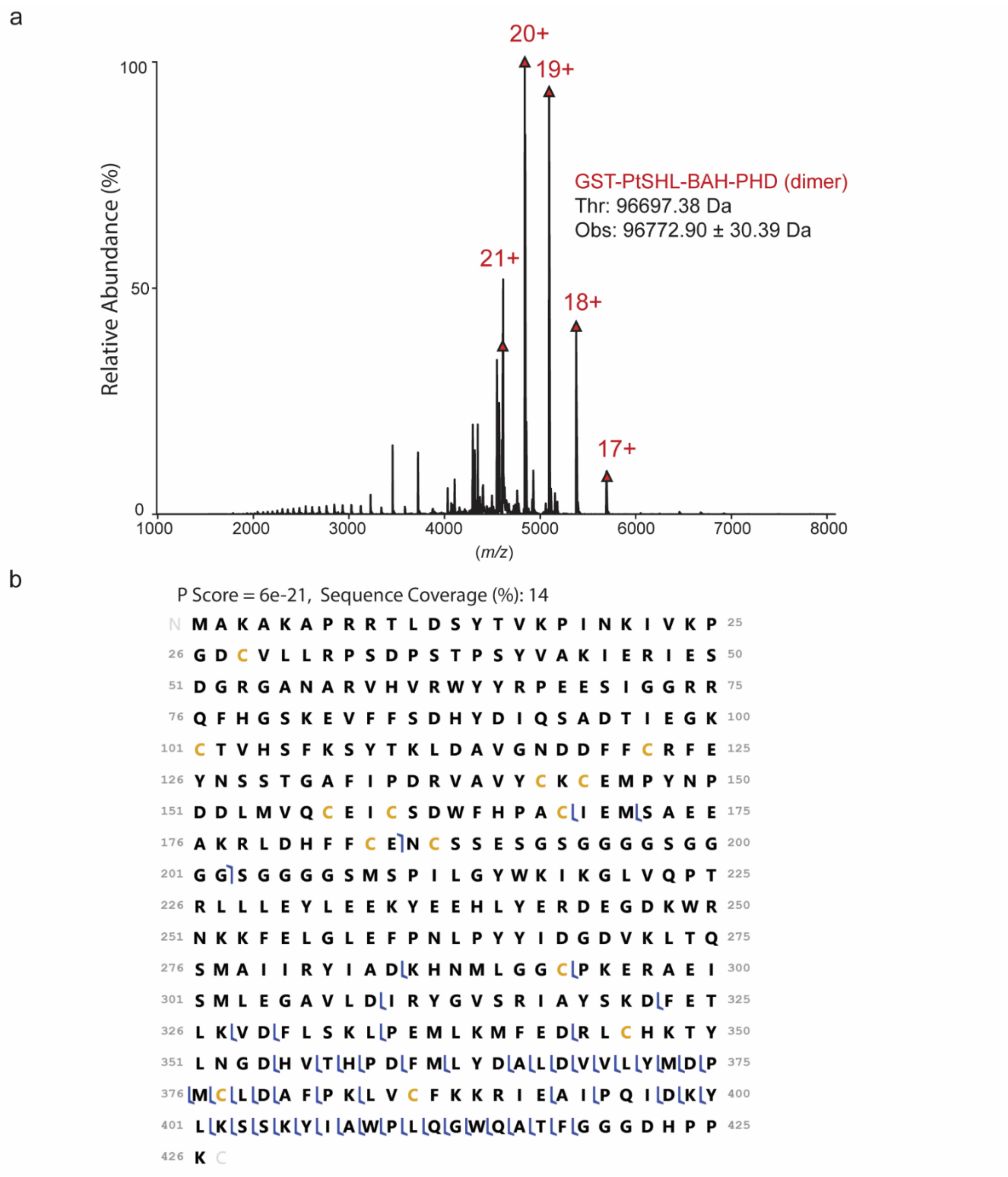


**SI Figure 15: Characterization of GST-PtSHL-BAH-PHD native tandem reader. a)** Representative Native MS1 analysis of GST-PtSHL-BAH-PHD dimer (possibly via GST-tag). Charge state distribution of intact GST-PtSHL-BAH-PHD dimer is represented in red (+17 to +21). **b)** Representative graphical fragment map of isolated GST-PtSHL-BAH-PHD fragmented by higher-energy collisional dissociation (HCD). Forward and reverse blue flags respectively represent *b* and *y* ions. Experiments were conducted with three independent biological replicates. Fragments were manually validated using TDValidator and corresponding P-scores were calculated using ProSight Lite.

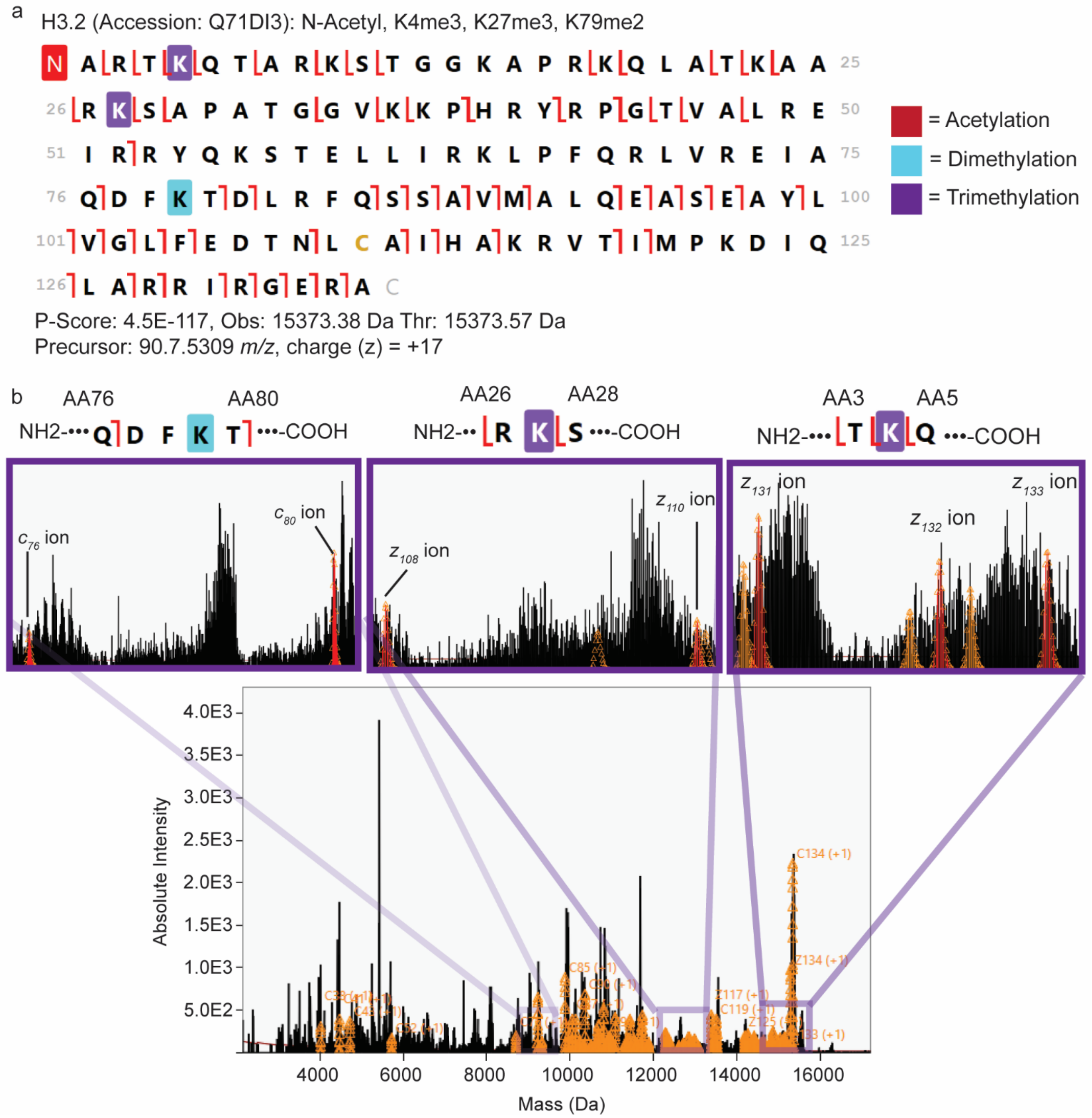


**SI Figure 16: Characterization of {H3.2 N-Acetyl, K4me3, K27me3, K79me2} proteoform from PtSHL-enriched endogenous nucleosomes. a)** Representative graphical fragment map of tandem MS fragmentation of {H3.2 N-Acetyl, K4me3, K27me3, K79me2} proteoform. Tandem MS fragmentation was performed using electron-transfer dissociation (ETD). Forward and reverse red flags respectively represent *c* and *z* ions. **b)** Deconvoluted mass spectra (Da) of MS/MS fragments of {H3.2 N-Acetyl, K4me3, K27me3, K79me2} proteoform. Orange represents matched fragment ions. Insets represent fragment ions in red localizing to K4me3, K27me3, and K79me2. Measurements were performed with three independent biological replicates. Observed (Obs) and theoretical (Thr) are represented as monoisotopic masses (Da). Fragments were manually validated using TDValidator and corresponding P-scores were calculated using ProSight Lite.

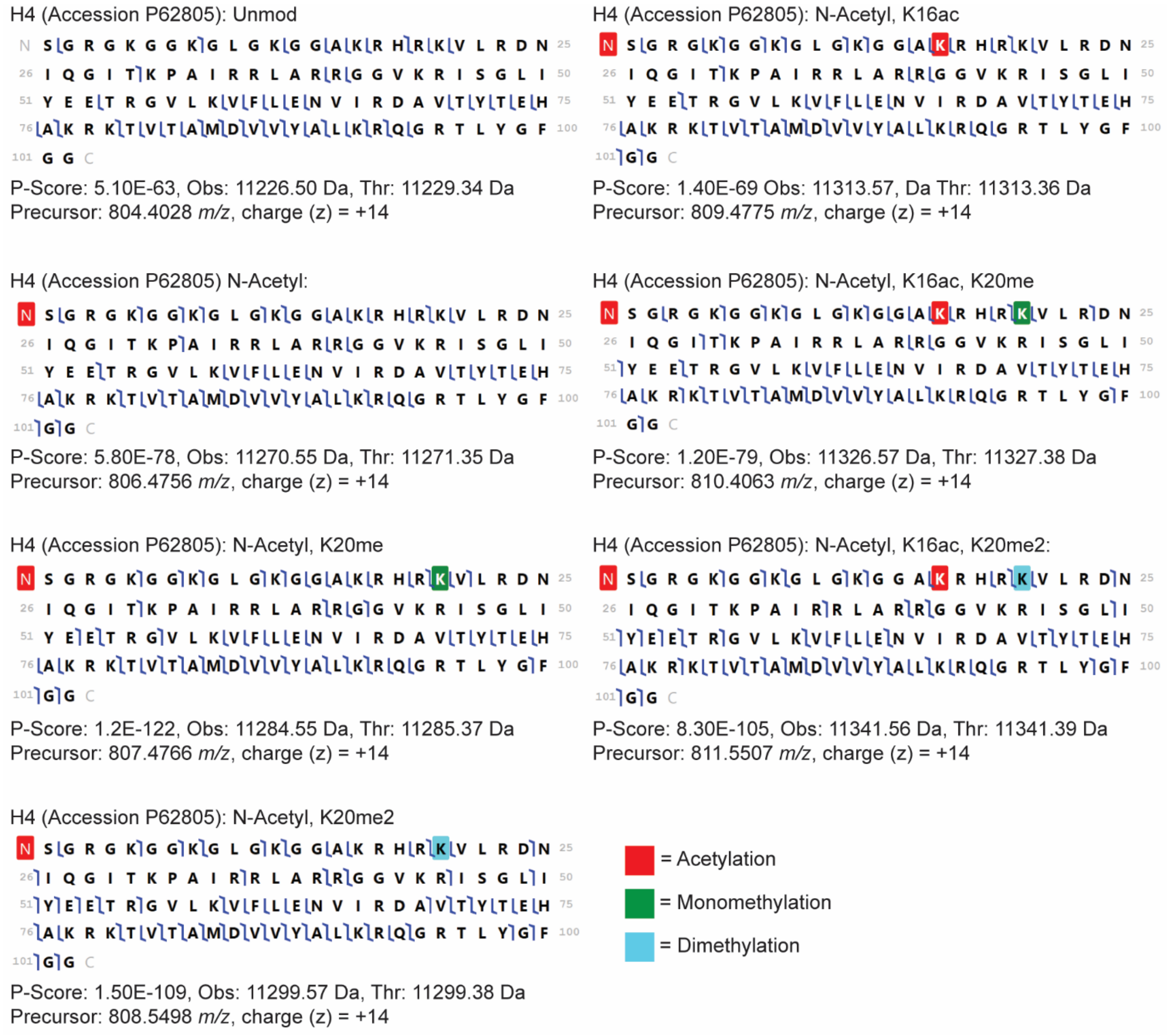


**SI Figure 17: Characterization of Histone H4 proteoforms from PtSHL-enriched endogenous nucleosomes.** Representative graphical fragment maps of tandem MS fragmentation of histone H4 proteoforms in PtSHL-enriched nucleosomes. Tandem MS fragmentation was performed using higher-collisional energy dissociation (HCD) across a distribution of H4 proteoforms (precursor ions (*m/z*) at charge state (z) +14 represent the intact H4 proteoforms used for tandem MS). Forward and reverse blue flags respectively represent *b* and *y* ions. Measurements were performed with three independent biological replicates. Observed (Obs) and theoretical (Thr) are represented as monoisotopic masses (Da). Fragments were manually validated using TDValidator and corresponding P-scores were calculated using ProSight Lite.

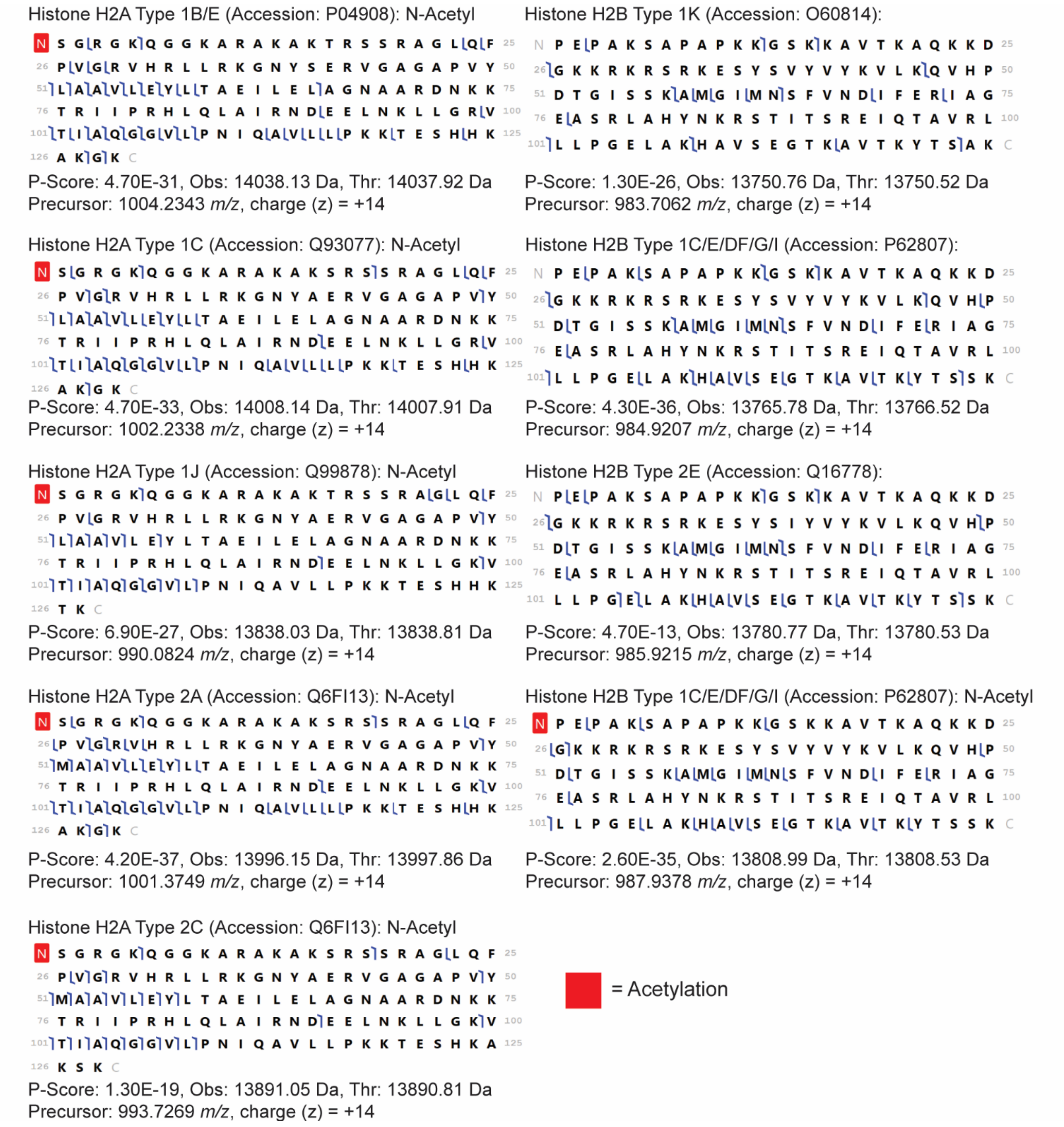


**SI Figure 18: Characterization of Histone H2A and H2B proteoforms from PtSHL-enriched endogenous nucleosomes.** Representative graphical fragment maps of tandem MS fragmentation of histone H2A and H2B proteoforms in PtSHL-enriched nucleosomes. Tandem MS fragmentation was performed using higher-collisional energy dissociation (HCD) across a distribution of H2A and H2B proteoforms (precursor ions (*m/z*) at charge state (z) +14 represent the intact H2A and H2B proteoforms used for tandem MS). Forward and reverse blue flags respectively represent *b* and *y* ions. Measurements were performed with three independent biological replicates. Observed (Obs) and theoretical (Thr) are represented as monoisotopic masses (Da). Fragments were manually validated using TDValidator and corresponding P-scores were calculated using ProSight Lite.

**SI Table 1A:** Relative Effective Concentration ($EC_{50}^{Rel}$) in Luminex assays to measure the interaction between GST-BPTF-PHD-BD constructs and PTM-defined histone H3 peptides.

| Construct | H3 1-20 peptide | | | H3K4me3 | | | H3 tetra$^{Ac}$ (K4ac, K9ac, K14ac, K18ac) | | | H3K4me3, tri$^{Ac}$ (K9ac, K14ac, K18ac) | | |
|---|---|---|---|---|---|---|---|---|---|---|---|---|
| | 1 | 2 | 3 | 1 | 2 | 3 | 1 | 2 | 3 | 1 | 2 | 3 |
| GST-BPTF-PHD-BD | 75 | 56 | 83 | 16.6 | 16.6 | 23.4 | 60.1 | 56.4 | 76.7 | 3.2 | 3.3 | 6.4 |
| GST-BPTF-PHD*-BD | 267 | 177 | 271 | 419 | 313 | 333 | 533 | 397 | 491 | 238 | 161 | 235 |
| GST-BPTF-PHD-BD* | 169 | 133 | 138 | 18.8 | 29.5 | 16.6 | 553 | 508 | 307 | 8.1 | 13.7 | 9 |
| GST-BPTF PHD*-BD* | 118 | 88 | 116 | 442 | 391 | 262 | 639 | 554 | 311 | 416 | 390 | 276 |

**SI Table 1B:** Averaged (AVG) and Standard Deviation (ST.DEV) of Relative Effective Concentration ($EC_{50}^{Rel}$) in Luminex assays to measure the interaction between GST-BPTF-PHD-BD constructs and PTM-defined histone H3 peptides.

| Construct | H3 1-20 peptide | | H3K4me3 | | H3 tetra$^{Ac}$ (K4ac, K9ac, K14ac, K18ac) | | H3K4me3, tri$^{Ac}$ (K9ac, K14ac, K18ac) | |
|---|---|---|---|---|---|---|---|---|
| | AVG | ST.DEV | AVG | ST.DEV | AVG | ST.DEV | AVG | ST.DEV |
| GST-BPTF-PHD-BD | 71.3 | 13.9 | 18.9 | 3.9 | 64.4 | 10.8 | 4.3 | 1.8 |
| GST-BPTF-PHD*-BD | 238.3 | 53.2 | 355.0 | 56.3 | 473.7 | 69.6 | 211.3 | 43.6 |
| GST-BPTF-PHD-BD* | 146.7 | 19.5 | 21.6 | 6.9 | 456.0 | 131.0 | 10.3 | 3.0 |
| GST-BPTF PHD*-BD* | 107.3 | 16.8 | 365.0 | 92.8 | 501.3 | 170.2 | 360.7 | 74.5 |

**SI Table 1C:** Relative Effective Concentration ($EC_{50}^{Rel}$) from Luminex assay between GST-tagged BPTF constructs and PTM-defined semi-synthetic nucleosomes.

| Construct | Unmodified Nuc | | | H3K4me3 | | | H3 tetra$^{Ac}$ (K4ac, K9ac, K14ac, K18ac) | | | H3K4me3, tri$^{Ac}$ (K9ac, K14ac, K18ac) | | |
|---|---|---|---|---|---|---|---|---|---|---|---|---|
| | 1 | 2 | 3 | 1 | 2 | 3 | 1 | 2 | 3 | 1 | 2 | 3 |
| GST-BPTF-PHD-BD | >1000 | 993 | >1000 | 128 | 85 | 123 | 324 | 240 | 378 | 25 | 16.4 | 28.7 |
| GST-BPTF-PHD*-BD | 762 | 671 | 738 | >1000 | >1000 | >1000 | >1000 | >1000 | >1000 | 340 | 300 | 382 |
| GST-BPTF-PHD-BD* | >1000 | >1000 | 776 | 356 | 509 | 234 | >1000 | >1000 | >1000 | 51 | 73 | 62 |
| GST-BPTF PHD*-BD* | 225 | 192 | 212 | 587 | 481 | 309 | 778 | 646 | 428 | 202 | 167 | 177 |

**SI Table 1D:** Averaged (AVG) and Standard Deviation (ST.DEV) of relative Effective Concentration ($EC_{50}^{Rel}$) from Luminex assay between GST-tagged BPTF constructs and PTM-defined semi-synthetic nucleosomes.

| Construct | Unmodified Nuc | | H3K4me3 | | H3 tetra$^{Ac}$ (K4ac, K9ac, K14ac, K18ac) | | H3K4me3, tri$^{Ac}$ (K9ac, K14ac, K18ac) | |
|---|---|---|---|---|---|---|---|---|
| | AVG | ST.DEV | AVG | ST.DEV | AVG | ST.DEV | AVG | ST.DEV |
| GST-BPTF-PHD-BD | >1000 | ND | 112.0 | 23.5 | 314.0 | 69.5 | 23.4 | 6.3 |
| GST-BPTF-PHD*-BD | 723.7 | 47.2 | >1000 | ND | >1000 | ND | 340.7 | 41.0 |
| GST-BPTF-PHD-BD* | >1000 | ND | 366.3 | 137.8 | >1000 | ND | 62.0 | 11.0 |
| GST-BPTF PHD*-BD* | 209.7 | 16.6 | 459.0 | 140.3 | 617.3 | 176.8 | 182.0 | 18.0 |

**Nuc-MS parameters**

- The Orbitrap Q Exactive Ultra High Mass Range (UHMR) and Orbitrap Ascend Tribrid MS are capable of scan ranges upwards to 8000 *m/z*. These MS instruments are most suitable for Nuc-MS analyses of readers and nucleosomes.
- The Orbitrap Ascend or Eclipse Tribrid MS was used specifically for the characterization of nucleosome-CAP complexes and their proteoforms due to multi-modal fragmentation approaches including Higher Collisional Dissociation (HCD), and Electron-Transfer Dissociation (ETD) allowing for increase in sequence coverage and PTM localization. **See Tables S3-S7.**

**SI Table 2:** Settings used for intact nMS analysis of BPTF:nucleosome complexes (MS1).

| | |
|---|---|
| **MS instrument** | Orbitrap Q Exactive UHMR MS |
| **Ion source type** | NSI |
| **Positive ion spray voltage (V)** | 1800-2500 |
| **Ion transfer tube temp (°C)** | 310 |
| **Pressure setting** | High Pressure Mode |
| **Scan type** | MS |
| **Detector type** | Orbitrap |
| **Orbitrap resolution at 400 *m/z*** | 6250 |
| **Mass range** | High |
| **Scan range (*m/z*)** | 500-15000 |
| **Microscan** | 2-20 |
| **RF lens (%)** | 150 |
| **AGC target** | $1x10^6$ |
| **Maximum injection time (ms)** | 20-175 |
| **Source fragmentation (V)** | 0 |
| **In-source trapping (IST) desolvation (V)** | -100 |
| **In-source trapping time (ms)** | 4 |

**SI Table 3:** Settings used for intact nMS analysis of CAPs (MS1).

| | |
|---|---|
| **MS instrument** | Orbitrap Ascend Tribrid MS |
| **Application mode** | Intact Protein |
| **Ion source type** | NSI |
| **Positive ion spray voltage (V)** | 1400-1800 |
| **Ion transfer tube temp (°C)** | 320 |
| **Pressure setting** | High Pressure Mode |
| **Scan type** | MS |
| **Detector type** | Orbitrap |
| **Orbitrap resolution** | 7500 |
| **Mass range** | High |
| **Scan range (*m/z*)** | 500-8000 |
| **Microscan** | 2 |
| **RF lens (%)** | 150 |
| **Normalized AGC target (%)** | 150 |
| **Maximum injection time (ms)** | 300 |
| **Source fragmentation (V)** | 40-80 |

**SI Table 4:** Settings used for MS/MS fragmentation analysis of CAPs (MS2).

| | |
|---|---|
| **MS instrument** | Orbitrap Ascend Tribrid MS |
| **Application mode** | Intact Protein |
| **Ion source type** | NSI |
| **Positive ion spray voltage (V)** | 1400-1800 |
| **Ion transfer tube ttemp (°C)** | 320 |
| **Pressure setting** | High Pressure Mode |
| **Scan type** | $MS^2$ |
| **Isolation width (m/z)** | BRD4: 500 *m/z*, DNMT3A-MPP8: 1000 *m/z*, PtSHL: 50 *m/z* |
| **Activation type** | HCD |
| **Collision energy type** | Normalized |
| **HCD collision energy (%)** | 20-55 |
| **Detector** | Orbitrap |
| **Orbitrap resolution** | 120000 |
| **Mass range** | High |
| **Scan range (*m/z*)** | 500-8000 |
| **RF lens (%)** | 150 |
| **Normalized AGC target (%)** | 1000 |
| **Maximum injection time (ms)** | 1000 |
| **Microscans** | 2 |
| **Source fragmentation (V)** | 40-80 |

**SI Table 5:** Settings used for characterization of ejected intact histone proteoforms from CAP-nucleosome complexes (MS1).

| | |
|---|---|
| **MS instrument** | Orbitrap Ascend Tribrid MS |
| **Application mode** | Intact Protein |
| **Ion source type** | NSI |
| **Positive ion spray voltage (V)** | 1400-1800 |
| **Ion transfer tube temp (°C)** | 320 |
| **Pressure setting** | Low Pressure Mode |
| **Scan type** | MS |
| **Detector type** | Orbitrap |
| **Orbitrap resolution** | 60000-120000 |
| **Mass range** | High |
| **Scan range (*m/z*)** | 500-8000 |
| **Microscan** | 1-2 |
| **RF lens (%)** | 150 |
| **Normalized AGC target (%)** | 150 |
| **Maximum injection time (ms)** | 500 |
| **Source fragmentation (V)** | 250 |

**SI Table 6:** Settings used for MS/MS fragmentation analysis of histone proteoforms (MS2).

| | |
|---|---|
| **MS instrument** | Orbitrap Ascend Tribrid MS |
| **Application mode** | Intact Protein |
| **Ion source type** | NSI |
| **Positive ion spray voltage (V)** | 1400-1800 |
| **Ion transfer tube temp (°C)** | 320 |
| **Pressure setting** | Low Pressure Mode |
| **Scan type** | $MS^2$ |
| **Isolation width (*m/z*)** | H2A: 20-30 *m/z*, H2B: 20-30 *m/z*, H3.2: 3.3 *m/z*, H4: 20-30 *m/z* |
| **Activation type** | HCD |
| **Collision energy type** | Normalized |
| **HCD collision energy (%)** | 30-60 |
| **Detector** | Orbitrap |
| **Orbitrap resolution** | 120000 |
| **Mass range** | High |
| **Scan range (m/z)** | 500-8000 |
| **RF lens (%)** | 150 |
| **Normalized AGC target (%)** | 1000 |
| **Maximum injection time (ms)** | 1000 |
| **Microscans** | 1-2 |
| **Source fragmentation (V)** | 250 |

**SI Table 7:** Settings used for Individual Ion Mass Spectrometry analysis of histone H3.2 proteoforms.

| | |
|---|---|
| **MS instrument** | Orbitrap Eclipse Tribrid MS |
| **Application mode** | Intact Protein |
| **Ion source type** | NSI |
| **Positive ion spray voltage (V)** | 2600 |
| **Ion transfer tube temp (°C)** | 320 |
| **Pressure setting** | Low Pressure Mode |
| **Scan type** | $MS^2$ |
| **Precursor (*m/z*)** | 904.7 |
| **Isolation width (m/z)** | 3.3 |
| **Activation type** | ETD |
| **ETD reaction time (ms)** | 3 |
| **ETD reagent target** | 8.0e4-1.0e5 |
| **Max ETD reagent injection time (ms)** | 200 |
| **Detector** | Orbitrap |
| **Orbitrap resolution** | 120000 |
| **Mass range** | High |
| **Scan range (*m/z*)** | 500-8000 |
| **RF lens (%)** | 150 |
| **Normalized AGC target (%)** | 5000 |
| **Maximum injection time (ms)** | 1000 |
| **Microscans** | 1 |
| **Source fragmentation (V)** | 250 |
